# Annelids win again: the first evidence of Hox antisense transcription in Spiralia

**DOI:** 10.1101/2021.01.30.428931

**Authors:** Elena L. Novikova, Nadezhda I. Bakalenko, Milana A. Kulakova

**Affiliations:** St. Petersburg State University, St. Petersburg, 199034 Russia

**Author notes:** Corresponding authors: ELN MAK.

**Keywords:** Annelids, Antisense RNA, Hox genes, lncRNA, Lophotrochozoa, NATs

## Abstract

To date it is becoming more and more obvious that multiple non-coding RNAs, once considered to be transcriptional noise, play a huge role in gene regulation during animal ontogenesis. Hox genes are key regulators of embryonic development, growth and regeneration of all bilaterian animals. It was shown that mammalian Hox loci are transcribed in both directions and noncoding RNAs maintain and control the normal functioning of Hox clusters. We revealed antisense transcripts of most of Hox genes in two lophotrochozoans, errant annelids *Alitta virens* and *Platynereis dumerilii.* It is for the first time when non-coding RNAs associated with Hox genes are found in spiralian animals. All these asRNAs can be referred to as natural antisense transcripts (NATs). We analyzed the expression of all detected NATs using sense probes to their Hox mRNAs during larval and postlarval development and regeneration by whole mount in situ hybridization (WMISH). We managed to clone several asRNAs *(Avi-antiHox4-1, Avi-antiHox4-2* and *Avi-antiHox5)* of these annelids and analyzed their expression patterns as well. Our data indicate variable and complicated interplay between sense and antisense Hox transcripts during development and growth of two annelids. The presence of Hox antisense transcription in the representatives of different bilaterian clades (mammals, myriapods and annelids) and similar expression relationships in sense-antisense pairs suggest that this can be the ancestral feature of Hox cluster regulation.

## Introduction

Eukaryotic genome mostly consists of non-coding sequences. About two thirds of the mammalian genome is actively transcribed but less than 2% of genomic DNA is protein-coding (Taft et al., 2007; Djebali et al., 2012). Among the plethora of non-coding regulatory transcripts such as microRNAs, PIWI-interacting RNAs, small nucleolar RNAs, and small interfering RNAs (Horabin, 2013; reviewed in Hombach and Kretz, 2016), an essential part is occupied by long non-coding RNAs (lncRNA). They are defined as non-coding transcripts that are more than 200 nt long and lack a long open reading frame (Brosnan and Voinnet, 2009). LncRNAs contain fewer exons than mRNAs and demonstrate a low expression level across the tissues (Derrien et al., 2012). Most of the known lncRNAs are transcribed with the help of the same transcription machinery as mRNAs, i.e. RNA-polymerase II, have a 5’ terminal methylguanosine cap and are generally spliced and polyadenylated (Ponting and Belgard, 2010). According to their position relative to protein-coding genes, lncRNAs can be intergenic (lincRNAs), intronic, that is, coded in gene introns, enhancer-associated (eRNAs) or transcribed from the same promoter as a protein-coding gene (pancRNA). They can also be transcribed from the strand lying opposite to the protein-coding one and overlapping with the coding sequence (He et al., 2008; Ponting et al., 2009; reviewed in Schmitz et al., 2016). The latter group of lncRNAs is referred to as Natural Antisense Transcripts (NATs) (Moran et al., 2012). According to some estimations, up to 70% of mammalian protein-coding genes possess natural antisense counterparts (Chen et al., 2004; Katayama et al., 2005; Faghihi and Wahlestedt, 2009).

LncRNAs participate in the regulation of multiple developmental processes such as pausing and release of gene transcription as an adaptation to heat shock during segmentation in *Drosophila* (Wang et al., 2007), stress response in *Arabidopsis thaliana* (Liu et al., 2012), sperm formation and the establishment of male identity in *Caenorhabditis elegans* (Nam and Bartel, 2012), stem cell fate regulation (Guttman et al., 2011), dose compensation (Meller and Rattner, 2002, Froberg et al., 2013) and cardiac development and heart function in mammals (Klattenhoff et al., 2013; Grote and Herrmann, 2013). An evidence of the dramatic role of lncRNAs in cancer and metastasis progression has been emerging in recent years (Gupta et al, 2010; Tang et al., 2016; Xie et al., 2016; Leng et al., 2018; Qian et al., 2018; Ghaforui-Fard et al., 2019).

The participation of lncRNAs in the control of Hox gene expression during embryonic and adult animal life has been shown in multiple works (Petruk et al., 2006; Rinn et al., 2007; Janssen and Budd, 2010; Wang et al., 2011; Gummalla et al., 2012). The tiling array of Hox genes from human fibroblasts representing 11 distinct positional identities revealed the transcription of 231 non-coding RNAs in four Hox loci (Rinn et al., 2007). There is evidence that most of these RNAs are transcribed from the strand opposite to Hox genes. It was shown that 48% of Hox lncRNAs displayed an expression pattern along the developmental axis and possessed the sequence motifs specific of ncRNAs located in distal, proximal or posterior sites of the human body (Rinn et al., 2007).

Though lncRNAs are abundant in Hox clusters of mammals, the characteristics and function of only a few of them have been described. Among them there are three lncRNAs from HoxA cluster: HOTAIRM1 (HOX antisense intergenic RNA myeloid 1), which is located between human *HOXA1* and *HOXA2* genes and transcribed antisense to HOXA genes (Zhang et al., 2009), *HOXA-AS2* localized between *HOXA3* and *HOXA4* (Zhao et al., 2013) and lincRNA *HOTTIP* localized upstream from *HOXA13* (Pradeepa et al., 2017). Long non-coding RNA *HOTAIR* (HOX Antisense Intergenic RNA) is localized between *HOXC11* and *HOXC12* genes and participates in the trans-regulation of genes from HOXD locus (Li et al., 2013). All the lncRNAs mentioned above are known to regulate various oncogenic processes such as breast cancer, esophageal cancer, lung cancer and gastric cancer (reviewed in Yu and Li, 2015; reviewed in Botti et al., 2018). A study of caecum development in herbivores and omnivores revealed the existence of two non-coding RNAs, named Hotdog (Hog) and Twin of Hotdog (Tog), which are located in a gene desert flanking the HoxD cluster. These RNAs share a common transcription start site and are specifically transcribed from opposite DNA strands in the growing caecum in midgestation (Delpretti et al., 2013).

It has been suggested that an intensive antisense and polycistronic transcription across mammalian Hox clusters may be the reason why Hox genes are packed close together in the mammalian clusters (Mainguy et al., 2007). These clusters are indeed relatively short and compact.

There are also several examples of antisense transcription within Hox clusters in the Ecdysozoan clade (Brena et al., 2006; Petruk et al., 2006; Janssen and Budd, 2010; Gummalla et al., 2012). Drosophila lncRNA *bxd* is transcribed from the area between *ultrabithorax* (*Ubx*) and *abdominal-A* (*Abd-A*) genes. It controls the *Ubx* expression through transcriptional interference, a mechanism ensuring down-regulation of 3’ genes by transcription of upstream ncRNAs through their promoters (Petruk et al., 2006; Pease et al., 2013). The same mechanism of gene regulation is at work in the 92-kb transcript *infraabdominal-8 (iab-8),* which is transcribed from the *iab* region localized between *Abd-A* and *Abd-B* genes (Gummalla et al., 2012). An antisense *Ubx* transcript is expressed in the development of myriapods *Strigamia maritima*, *Glomeris marginata* and *Lithobius forficatus* (Brena et al., 2006; Janssen and Budd, 2010).

The presence of antisense transcription within Hox clusters in the representatives of two different clades brings up a question of whether this is a plesiomorphic bilaterian feature or an invention of mammals serving to support the integrity of their compact Hox clusters. To answer this question, an evidence from the third clade, the Lophotrochozoa, is strongly needed. At present, there are no data about lncRNA Hox expression, particularly antisense RNA (asRNA) transcription, in this clade.

There are two options to detect antisense transcription by whole mount in situ hybridization (WMISH). In case the sequence of antisense transcript significantly overlap the sense one (by more than 500 bp) we can use sense probe to mRNA sequence to reveal antisense transcription. The second option is to clone the antisense fragment and use the probe (antisense probe) to this fragment to detect antisense transcription in WMISH. In this study, we revealed antisense transcripts of most of the Hox genes in two lophotrochozoans, errant polychaetes *Alitta virens* and *Platynereis dumerilii.* We managed to clone several asRNAs *(Avi-antiHox4-1, Avi-antiHox4-2* and *Avi-antiHox5)* of these annelids and analyzed their expression during larval and postlarval development and regeneration by WMISH.

## Results

### Cloning of sense and antisense Hox transcripts

Cloning of sense and antisense Hox transcripts of A. virens and P. dumerilii was mostly performed in the Laboratory for Development and Evolution, Department of Zoology, University of Cambridge). The search and cloning of rare low-abundant RNAs is technically complicated. We used the RACE method (Rapid Amplification of cDNA Ends) which allowed us to obtain 3’- and 5’-ends of target RNAs. Since we observed the localization of all antisense transcripts detected by sense probes to mRNA sequences in the cytoplasm we suggested that these transcripts are polyadenylated from 3’-end. Thus, we performed amplification of transcript 3’-end using the primer, which contained polyT sequence. This sequence is complementary to polyA on the 3’-end of target transcript. For the amplification of 5’-ends SMARTTM (Switching Mechanism At 5’ end of the RNA Transcript) RACE was used (Clontech).

For primer construction we used the known sequences of sense Hox gene fragments. We constructed three pairs of forward and reverse primers for different sites of each Hox sequence. The primers’ length varied from 23 to 28 nt with Tm around 70 ^0^C. The cloned sequences were ligated into pGEM®-T Easy Vector (Promega).

As a result the following antisense transcripts were cloned (Table S1):

1. a 3’ fragment of *Avi-antiHox5* (Gene Bank KP100547). This fragment is 938 nt long and complementary to the 5’-end of *Avi-Hox5* mRNA and 5’UTR.
2. a 3’ fragment of *Avi-antiHox4 (Avi-antiHox4_1;* Gene Bank KX998894.1). This fragment is 1212 nt long and complementary to the 3’-exon of *Avi-Hox4* mRNA (Fig. 1).
3. a 3’ fragment of *Avi-antiHox4 (Avi-antiHox4_2;* Gene Bank KX998895.1). This fragment is 739 nt long and includes a fragment of the 5’ intergenic region (Fig. 1).
4. a 5’ fragment of *Pdum-antiHox7* which is complementary to the 3’-end of *Pdum-Hox7* and 464 nt long.
5. a 5’ fragment of *Avi-antiHox7* which is complementary to the 3’-end of *Avi-Hox7* and 1244 nt long.

**Figure 1.**
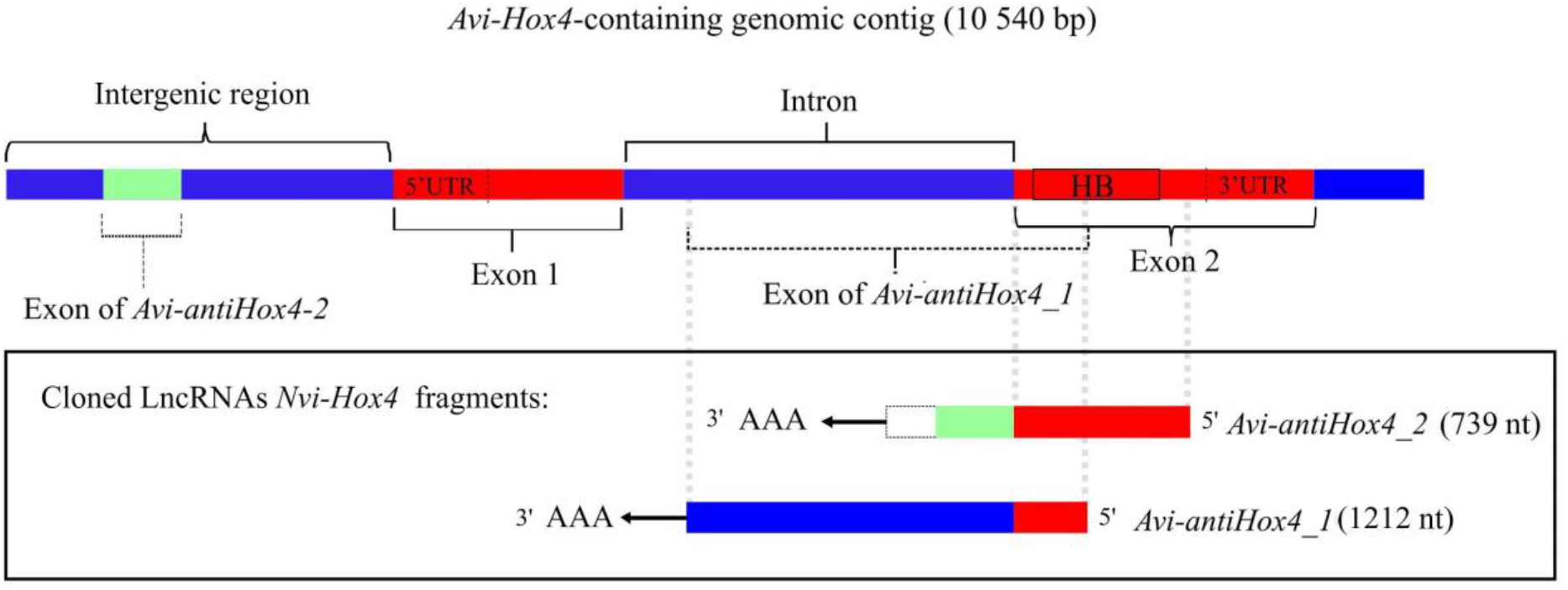
Schematic representation of *Avi-Hox4* sequence with the projection of antisense cloned fragments of *Avi-antiHox4_2* and *Avi-antiHox4_1*. Red color marks exons of the sense transcript. Blue color marks intergenic areas and introns, one of which is a part of *Avi-antiHox4_1* exon. Light green color marks the exon of *Avi-antiHox4_2*, which is localized upstream to the sequence that codes *Avi-Hox4* mRNA. White color indicates an exon of the *Avi-antiHox4_2* fragment, which was not found inside the known genomic sequence. The positions of introns and exons were obtained from genomic sequence locally assembled from short genomic reads using Geneious Prime software (data not shown).

### In situ hybridization

We have previously published a detailed description of the expression of coding transcripts from the *Avi-Hox* genes during development and regeneration of *A. virens* (Kulakova et al., 2007, Bakalenko et al., 2013, Novikova et al., 2013). Therefore, here we present these sense patterns only for the sake of comparison with the antisense expression patterns. Such comparisons are essential for understanding the transcription dynamics and the possible roles of the antisense transcripts.

#### A note on Terminology

We use the term “antisense RNA probe” to mean a probe containing the sequence which is antisense with respect to the coding strand of the Hox gene, and which therefore hybridises to and detects transcripts from the strand that encodes the Hox protein. Conversely, the term “sense RNA probe” is used to mean a probe that contains sequences from the sense (i.e. Hox protein encoding) sequence of the gene. Such probes will hybridise to antisense transcripts derived from the same genomic region.

#### antiHox1

To detect antisense transcription from the region of the *Avi-Hox1* gene, we used a digoxigenin-labelled *A. virens* sense RNA probe (548 bp) (Fig. 2 A) that overlapped the homeobox (HB) and 3’UTR of *Avi-Hox1. Avi-antiHox1* is expressed in segment ectoderm, the coelom and the gut in the posterior quarter of the body of juvenile and regenerating worms. Weak expression is visible in pygidial cirri and the esophageal region (Fig. 2 C (a, b, c); B, orange color). A probe containing the antisense strand of this same sequence, complementary to mRNA, clearly reveals sense transcription in the ganglia of young but already formed segments, in the cirri and in the esophageal area (Fig. 2 D (a, b, c); B, green color). It is noteworthy that the domain of maximal expression of *Avi-antiHox1* coincides with one of the zones of minimal *Avi-Hox1* transcription (Fig. 2 C (b), D (b)). However, there are areas (esophagus, pygidial cirri) where both transcripts are visible.

**Figure 2.**
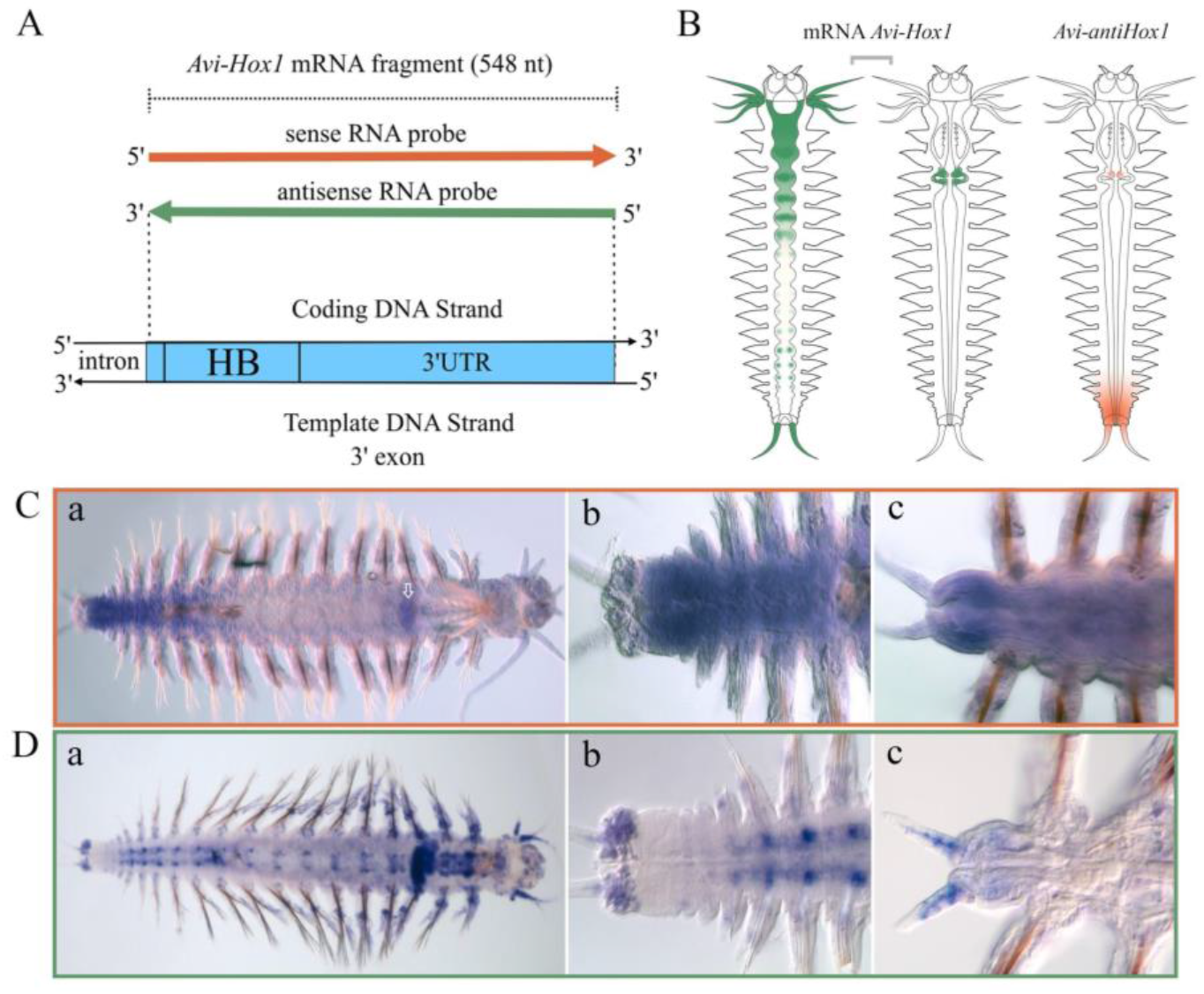
Sense and antisense transcription of the *Hox1* gene in juvenile *A. virens.* **A.** Schematic representation of the probes used with their projection onto the genomic sequence. **B.** Schematic representation of expression patterns for *Avi-Hox1* (sense) and *Avi-antiHox1* (antisense) transcripts (shown in green and orange respectively). **C.** WMISH with sense strand as probe. Expression of *Avi-antiHox1* RNA in juvenile worm (**a**, **b**) and regenerating worm at 3 days after caudal amputation (**c**) (orange framework); White outline arrow in (**a**) points to the localized signal in the esophagus. **D.** (**a**-**c**) WMISH with antisense strand as probe. Expression of *Avi-Hox1* mRNA in juvenile worm (**a**, **b**) and regenerating worm at 3 days after caudal amputation (**c**) (green framework). In **C** and **D**, the worms are oriented with their heads to the right on all photos.

#### antiHox2

Similarly to *antiHox1,* the transcription of antisense *Hox2* of *A. virens* was revealed with sense RNA probe (580 bp) synthesized from 3’-area of coding *Avi-Hox2* sequence (Fig. 3 A). *Avi-antiHox2* transcript is detected in the semicircle of the ectodermal cells on the dorsal site of prepygidial area and in the esophagus (Fig. 3 C (a, b, c); B, orange color). The first expression domain probably corresponds to the ectodermal growth zone. The second domain coincides with the territory of *Avi-antiHox1* expression (Fig. 2 C (a); B, orange color). *Avi-Hox2* mRNA transcripts are visible in the mesoderm of the youngest segments, presumably, in the mesodermal growth zone and at the base of the jaws (Fig. 3 D (a, b, c); B, green color). It is interesting that the expression zones of sense and antisense *Avi-Hox2* transcripts do not overlap but adjoin each other.

**Figure 3.**
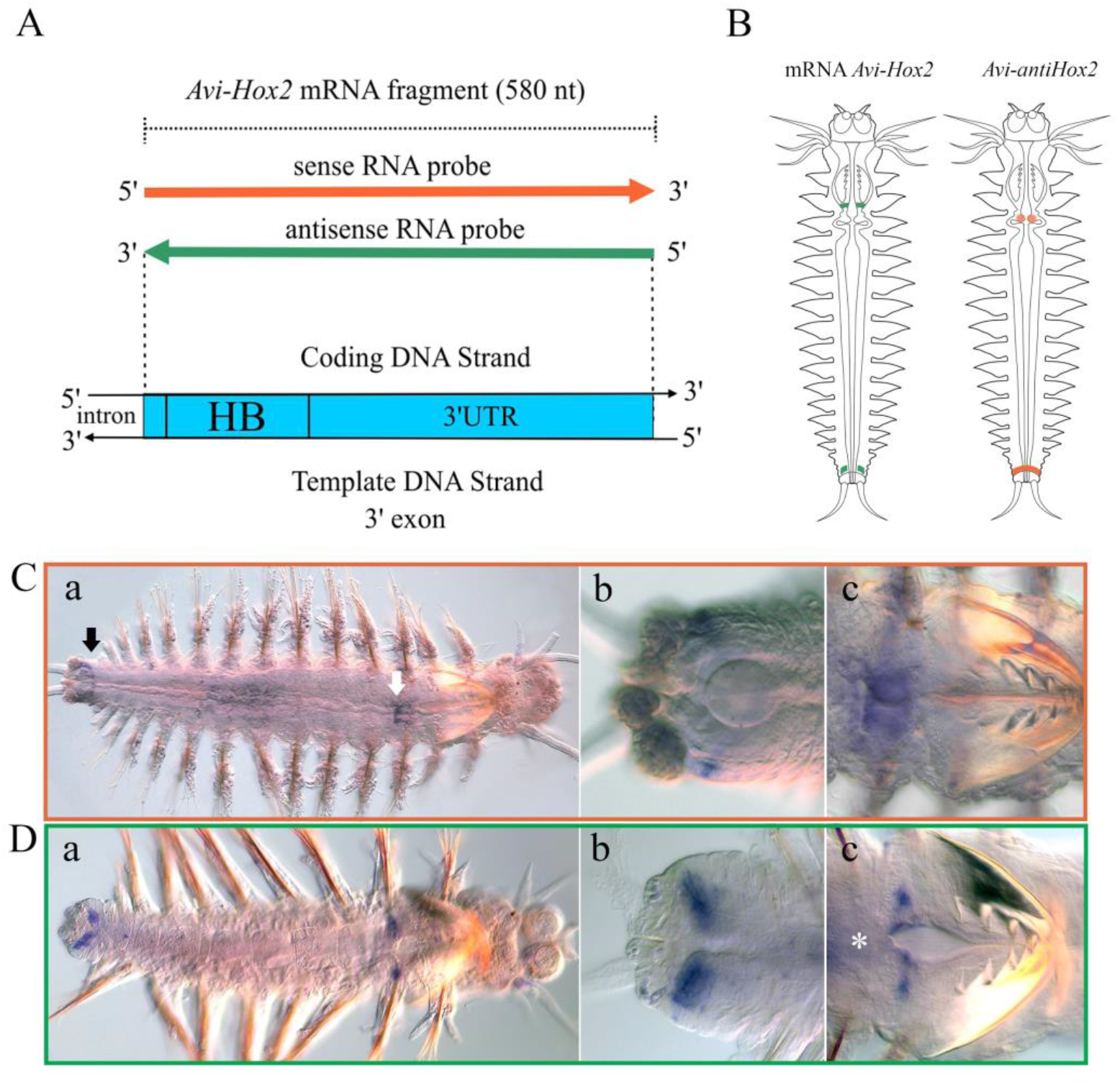
Sense and antisense transcription of the *Hox2* gene in juvenile *A. virens.* **A.** Schematic representation of the probes used with their projection onto the genomic sequence. **B.** Schematic representation of expression patterns for *Avi-Hox2* (sense) and *Avi-antiHox2* (antisense) transcripts (shown in green and orange respectively). **C.** WMISH with sense strand as probe detects *Avi-antiHox2* transcription in juvenile worms (**a**-**c**, orange framework). Back arrow (**a**) indicates dorsal semicircle of ectodermal cells with the signal; white arrow indicates the signal in esophagus; **b** - deep optical slice, demonstrating position of *Avi-antiHox2* RNA in the ectodermal part of the growth zone; (**c)** presents expression in esophagus. **D.** WMISH with antisense strand as probe detects *Avi-Hox2* transcription in juvenile worms (**a**-**c**, orange framework). (**b)** - deep optical slice, demonstrating position of *Avi-Hox2* mRNA in the mesodermal part of the growth zone; c presents the expression in the pharynx. The position of esophagus is marked by asterisk. In **C** and **D**, the worms are oriented with their heads to the right on all photos.

#### antiHox3

We failed to detect an antisense transcription of *A. virens Hox3* with the sense RNA probe (550 nt) corresponding to the second exon and 3’UTR of *Avi-Hox3* mRNA. However, a weak antisense transcription of *Hox3* is visible in *P. dumerilii*. To reveal it, we used sense RNA probe (619 nt) that overlaps the first exon without 5’UTR and partially the second exon (Fig. 4 A). *Pdum-antiHox3* expression is detected only at the nectochaete stage in the forming esophagus (Fig. 4 B (a)). We cannot be sure that antisense RNA of this gene is transcribed in such a narrow frame. This result can be due to the fact that we detect only the maximal expression, which occurs at the nectochaete stage. mRNA of *Pdum-Hox3* marks the territory of the future growth zone from the metatrochophore stage, and this expression is retained in juvenile worms (Fig. 4 B (b)).

**Figure 4.**
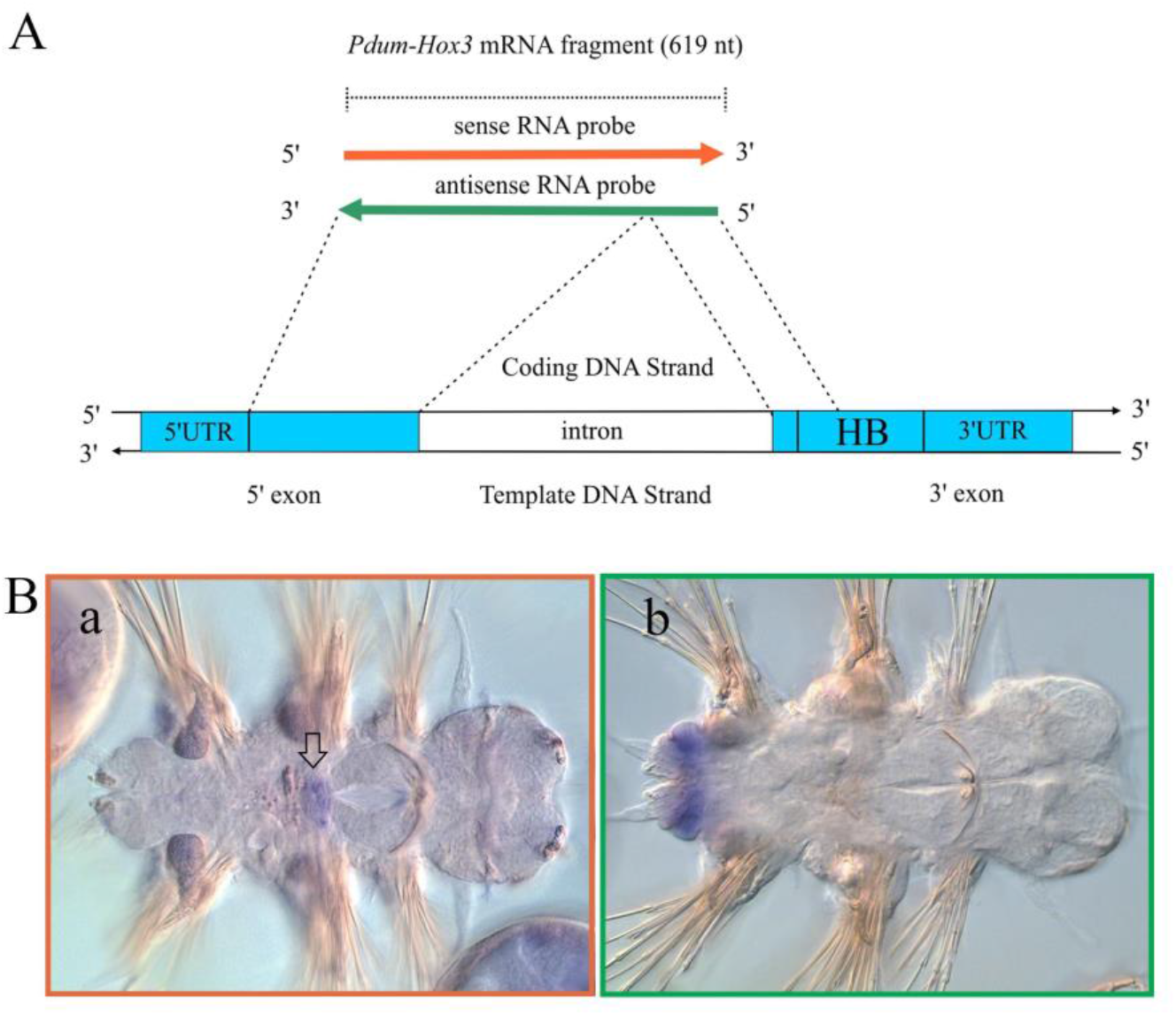
Sense and antisense transcription of the *Hox3* gene in *P. dumerilii* nectochaete. **A.** Schematic representation of the probes used with their projection onto the genomic sequence. **B.** WMISH with sense strand as probe detects *Pdum-antiHox3* transcription in esophagus anlagen (orange framework, black outline arrow) (**a**); WMISH with antisense strand as probe detects *Pdum-Hox3* transcription in the forming growth zone (green framework) (**b**). **B** - head to the right.

#### antiHox4

To analyze the antisense transcription of *A. virens Hox4*, we used an antisense RNA probe for the largest of the cloned fragments, *Avi-antiHox4_1* (1212 nt), and a sense RNA probe for 3’-fragment of *Avi-Hox4* mRNA (618 nt) (Fig. 5 A). *Avi-antiHox4_1* is detected for the first time at the early trochophore stage at 50 h. The expression domain is represented by paired dots in the nuclei of mesodermal band cells (Fig. 5 D (b)). In four hours the antisense RNA is revealed in the nuclei of adjacent ectodermal cells that will contribute to the formation of the third larval segment and the pygidium (Fig. 5 D (c)). Four hours later *Avi-antiHox4_1* transcription starts in the cytoplasm (Fig. 5 D (d)). The detected signal gradually loses the nuclear localization and by the late trochophore stage is retained only in the cytoplasm of the ectodermal cells (Fig. 5 D (e, f)). At the metatrochophore stage (120 h) the transcript is revealed in the area of the future pygidium and in a few surface cells, localized along the midline of elongating vegetal plate (Fig. 5 D (g)). *Avi-antiHox4_1* expression persists in the pygidium, pygidial cirri and ectodermal growth zone at the nectochaete stage (Fig. 5 D (h)).

**Figure 5.**
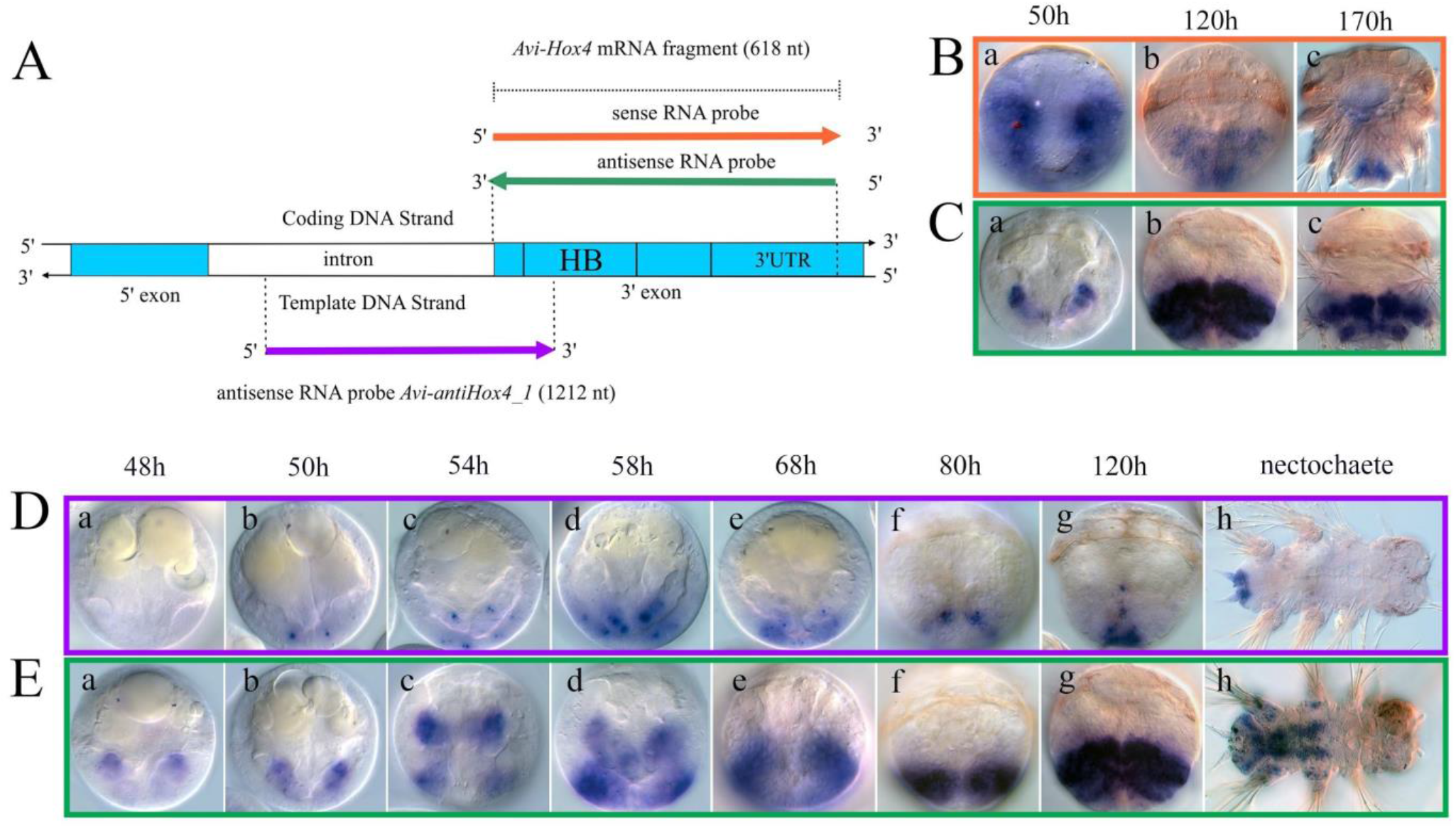
Sense and antisense transcription of the *Hox4* gene at various larval stages of *A. virens.* **A.** Schematic representation of the probes used with their projection onto the genomic sequence. Purple color indicates the probe to the cloned sequence *Avi-antiHox4_1*. **B.** WMISH with sense strand as probe detects *Avi-antiHox4* transcription at trochophore (**a**), metatrochophore (**b**) and nectochaete (**c**) stages (orange framework). **C.** WMISH with antisense strand as probe detects *Avi-Hox4* transcription at the same stages as in **B** (green framework). **D.** WMISH with the probe to *Avi-antiHox4_1* at trochophore (**a**-**f**), metatrochophore (**g**) and nectochaete (**h**) stages (purple framework). **E.** WMISH with antisense strand as probe detects *Avi-Hox4* transcription at the same stages as in **D** (**a**-**h**; green framework). The full description of the transcription pattern is given in the text. **B**, **C**, **D**, **E** - ventral view; episphere to the top; (**h)** - head to the right.

It is interesting to compare the distribution of *Avi-antiHox4_1* with the pattern of mRNA expression of *Avi-Hox4*. This gene works on the territory of the second and the third larval chaetiger with an early expression initiation at 46 hpf (hour post fertilization). Its transcript is detected in the mesodermal bands and larval ectoderm (Fig. 5 E (a-h)). Previously we described in detail the mesodermal expression of *A. virens* Hox genes (Kulakova et al., 2017). The sense transcript of *Avi-Hox4* appears four hours after the antisense one and demonstrates only cytoplasmic localization. Before the metatrochophore stage the territories of sense and antisense transcription partially overlap at the level of the posterior boundary of *Avi-Hox4* domain and the anterior boundary of *Avi-antiHox4_1* domain. Later the areas of their transcription overlap in pygidial cirri (Fig. 6 A (a, e)). We failed to detect the expression pattern of the smaller of the two cloned antisense transcripts *(Avi-antiHox4 2*, 739 nt) (Fig. 1).

**Figure 6.**
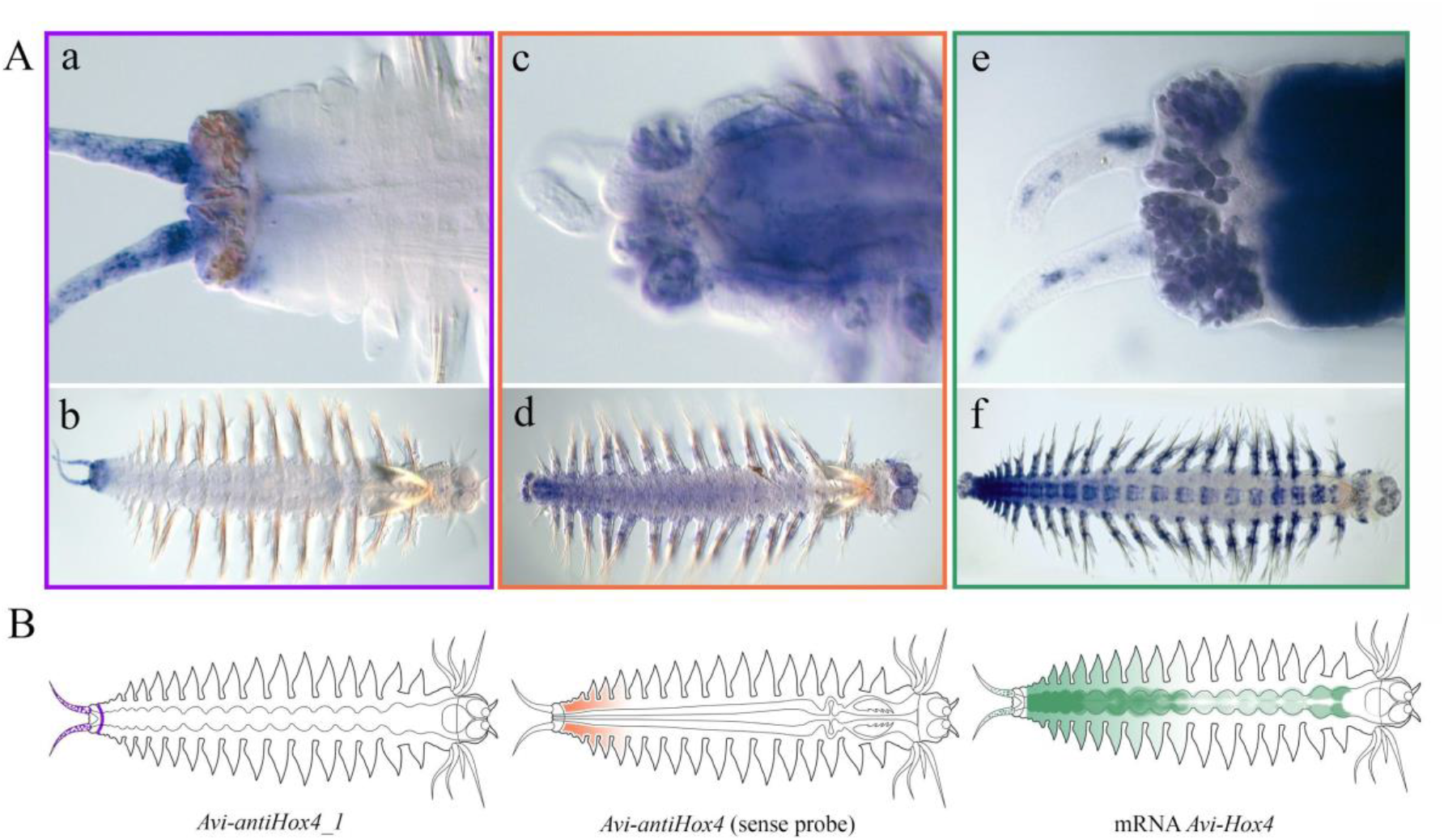
Sense and antisense transcription of the *Hox4* gene in juvenile *A. virens.* **A.** WMISH with the probe to *Avi-antiHox4_1* transcript (**a**, **b**; purple framework). WMISH with sense strand as probe detects 3’-associated *Avi-antiHox4* transcript (**c**, **d**; orange framework); WMISH with antisense strand as probe detects *Avi-Hox4* transcription (**e**, **f**; green framework); **B.** Schematic representation of expression patterns of all studied *Hox4* transcripts in juvenile worms. **A** (**a**, **c**, **e**) - heads to the right. The full description is in the text.

Using the sense RNA probe covering the second exon, we revealed one more antisense transcript with a different expression pattern than that described above. This transcript is detected at the trochophore stage at the margins of the closing vegetal plate (Fig. 5 B (a)). Noteworthy, there is no sense transcription in the ectoderm at this stage, and the cloned antisense transcript (*Avi-antiHox4_1)* is detected in the mesodermal cells (Fig. 5 D (b), E (b); C (a)). At the early metatrochophore stage the sense RNA probe is visible in the surface cells of the ventral part of the vegetal plate in all the three larval segments. Sense and antisense transcription domains partially overlap at this stage (Fig. 5 B (b), C (b)). Later, the ectodermal expression *Avi-antiHox4* (non-cloned) vanishes, and by the late metatrochophore stage the signal is detected only in deep, presumably mesodermal cells at the basis of the pygidial anlagen. During this period, the sense transcript is localized in the segmental ectoderm (Fig. 5 B (c), C (c)).

In juvenile worms *Avi-antiHox4_1* is detected only in the ectodermal growth zone and in the pygidium and cirri (Fig. 6 A (a, b); B, purple color). The non-cloned antisense transcript that was detected by the sense RNA probe is localized in the mesoderm of the young segments (Fig. 6 A (c, d); B, orange color). *Avi-Hox4* mRNA marks the ectoderm of young segments, the ganglia of the ventral nerve cord and individual cells in pygidial cirri (Fig. 6 A (e, f); B green color). Thus, in juvenile worms the sense transcript and both antisense ones are localized in a complementary manner in non-overlapping zones. The only area where sense and antisense transcripts seem to overlap is the pygidial cirri. However, since *Avi-antiHox4_1* demonstrates a non-homogeneous distribution and the sense RNA is detected in separate cells, we can assume that their localization in pygidial cirri is also complementary. Double WMISH is necessary to ascertain the co-localization of these transcripts.

Using the sense RNA probe (592 nt) (Fig. 7 A), we managed to detect an antisense transcript of *Hox4* gene of *P. dumerilii* (Fig. 7 B (a, b, c)). We determined the localization of this transcript in the ectoderm. At the trochophore stage its expression pattern partially overlaps with the transcription zone of the sense RNA (Fig. 7 B (a), C (a)), but the situation changes dramatically by the nectochaete stage. The sense transcript is clearly visible in the ectoderm of the second and the third segments, while the antisense transcript is detected in the pygidium, the stomodeum and in a few cells of the head (Fig. 7 B (b), C (b)).

**Figure 7.**
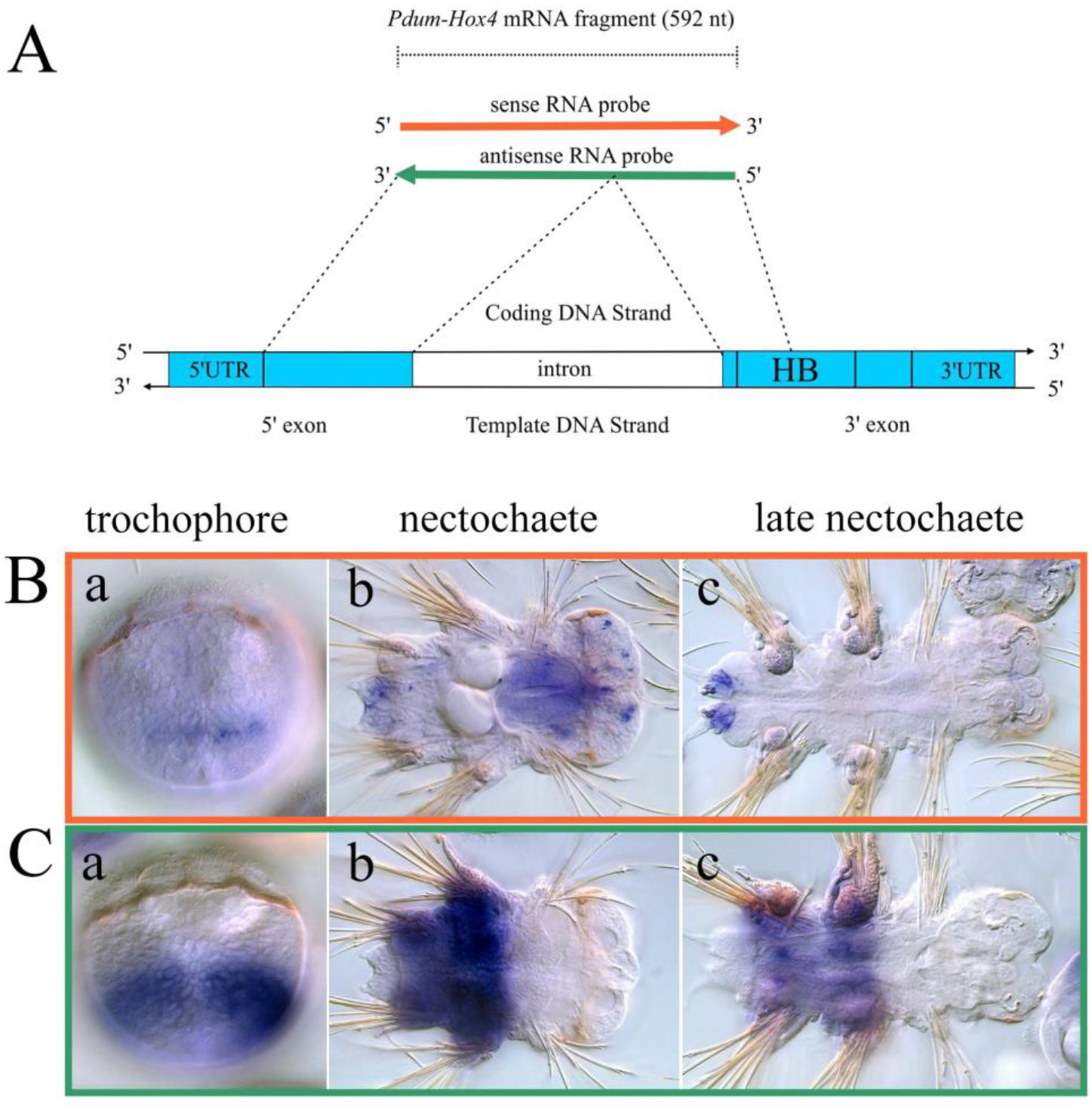
Sense and antisense transcription of the *Hox4* gene at the various larval stages of *P. dumerilii.* **A.** Schematic representation of the probes used with their projection onto the genomic sequence. **B.** WMISH with sense strand as probe detects *Pdum-antiHox4* transcript (**a**-**c**; orange framework). **C.** WMISH with antisense strand as probe detects *Pdum-Hox4* transcript (**a**-**c**; green framework). (**a**) - ventral view, episphere to the top. (**b**, **c**) - ventral view, head to the right. The full description is in the text.

Noteworthy, *Avi-Hox4* antisense RNA is not detected in the mouth and in the episphere (Fig. 5 D (h)). Later, *Pdum-antiHox4* is transcribed only in the pygidium, strictly outside the territory of *Pdum-Hox4* mRNA expression (Fig. 7 B (c), C (c)).

*Pdum-antiHox4* transcription pattern reminds in some respects that of *Avi-antiHox4_1,* but we cannot consider these transcripts as homologous. Firstly, *Pdum-antiHox4* is transcribed from the sequence of the first and the second exons, while *Avi-antiHox4_1* does not include the first exon (Fig. 1). Secondly, even if we assume that the *Pdum-Hox4* sense RNA probe reveals *Pdum-antiHox4* only at the level of the second exon, a methodological contradiction arises: the part of the probe including the second exon is too short (127 nt) for the transcript to be revealed by WMISH.

#### antiHox5

Previously we described the transcription pattern of *Avi-Hox5* asRNA revealed by the sense RNA probe (1030 nt) in juvenile worms during normal growth and regeneration (Bakalenko et al., 2013; Novikova et al., 2013). The cloned fragment of *Avi-Hox5* mRNA used for the synthesis of sense and antisense probes includes a part of the first exon and the complete proteincoding sequence of the second exon with a short 3’UTR area (Fig. 8 A). We managed to clone a fragment of asRNA of *Avi-Hox5* gene, which is complementary to 5’UTR of the coding transcript, and a small protein-coding area, which overlaps the activator domain (Fig. 8 A). We analyzed the transcription pattern of the cloned *Avi-antiHox5* (GenBank: KP100547.1) and compared it to the pattern of the protein-coding sequence and the sense RNA probe pattern.

**Figure 8.**
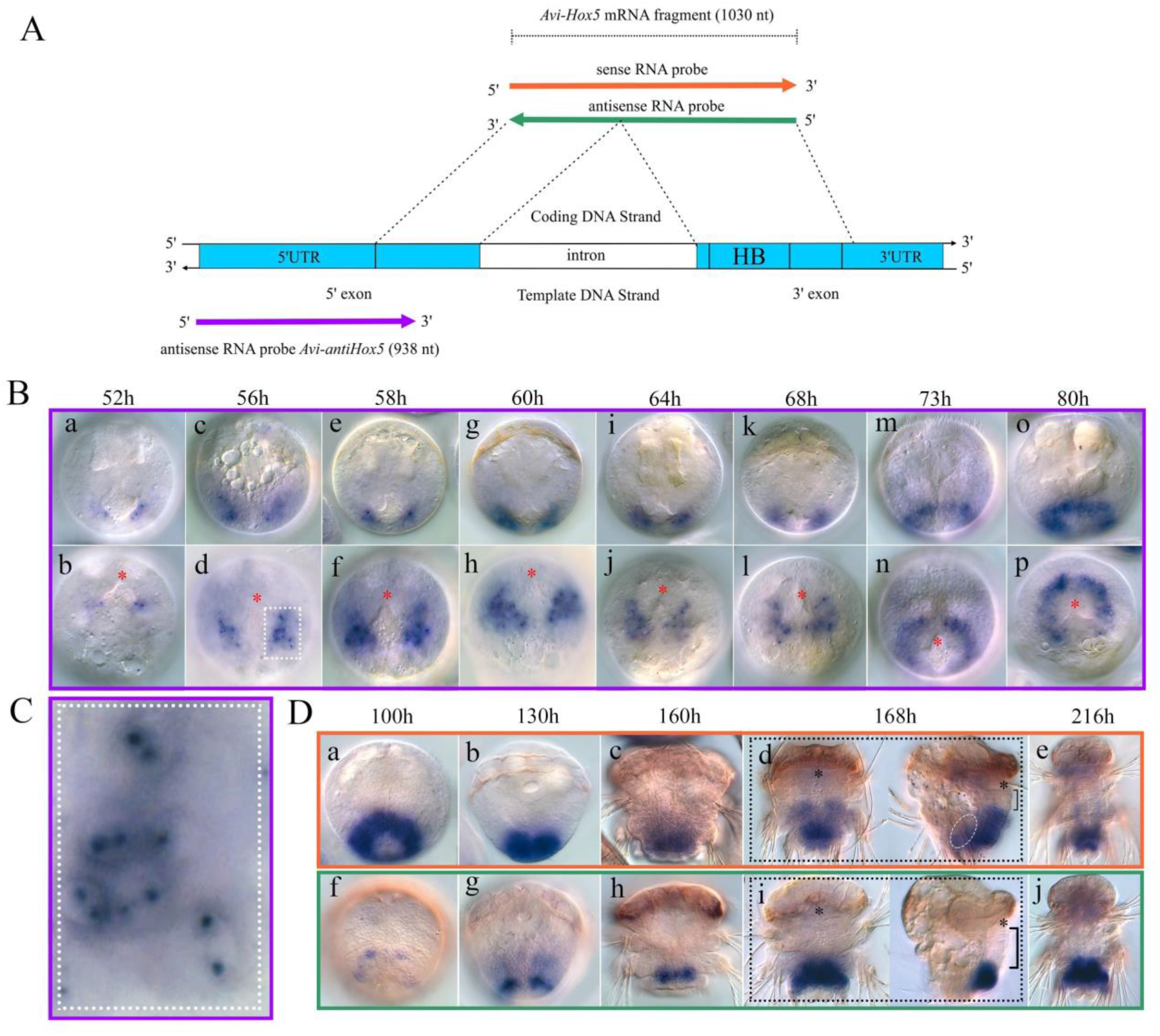
Sense and antisense transcription of the *Hox5* gene at the larval stages *A. virens*. **A.** Schematic representation of the probes used with their projection onto the genomic sequence. Purple color indicates the probe to the cloned sequence of *Avi-antiHox5*. **B.** WMISH with the probe to *Avi-antiHox5* transcript (**a**-**p**; purple framework). Early stage trochophore larvae (**a**, **c**, **e**, **g**) and middle stage trochophore larvae (**i**, **k**, **m**, **o**) are depicted in frontal projection (**a**, **c**, **e**, **g**, **i**, **k**, **m**, **o**) with the reducing depth of optical slices as the signal moves to the ventral side. In the bottom row the larvae are orientated with their vegetal pole to the front (**b**, **d**, **f**, **h**, **j**, **l**, **n**, **p**), ventral side is to the top. Red asterisk marks the vegetal pole. At **C** the enlarged fragment of **B** (**d**) is presented where the nuclear localization of the transcripts in two symmetrical domains is clearly visible. **D.** WMISH with sense strand as probe detects *Avi-antiHox5* transcript (orange framework) at metatrochophore (**a**, **b**) and nectochaete (**c**-**e**) stages; WMISH with antisense strand as probe detects *Avi-Hox5* transcript (green framework) at metatrochophore (**f**, **g**) and nectochaete (**h**-**j**) stages. On (**d**) and (**i**) the larvae of the same age are presented from the ventral side (left) and in lateral position (right, deep optical slice) to demonstrate the mesodermal localization of antisense transcript (white dotted line). Right square bracket marks the territory of the ventral neuroectoderm with no sense and antisense signal, the black asterisk marks the larval mouth. The full description of the transcription pattern is in the text.

Both probes to antisense RNAs demonstrate similar transcription dynamics and patterns. Here we describe the early expression of the cloned fragment because it was studied in more detail (Fig. 8 B (a-p), C). The antisense transcription starts before the sense one. *Avi-Hox5* mRNA is first detected at the late trochophore stage (Fig. 8 D (f)), while both antisense transcripts are first visible almost 50 hours earlier at the early trochophore stage (Fig. 8 B (a, b) (shown for cloned asRNA). The antisense expression starts in the bilaterally symmetrical groups of ectodermal cells localized close to the vegetal pole of the larva (Fig. 8 B (a - f)). Initially asRNA is detected only in the nuclei (52 hpf), entering the cytoplasm in about 6 hours (58 hpf) (Fig. 8 C). The number of positive cells rapidly increases, and they form a ring by the middle trochophore stage (60-80 hpf; Fig. 8 B (g - p)).

Noteworthy, *Avi-antiHox5* retains the nuclear localization starting from very early stages (52-56 hpf) and to the middle trochophore stage (80 hpf). Similar to *Avi-antiHox4_1* (Fig. 5 D (b, c)), this transcript forms paired conglomerates in the nucleus (Fig. 8 C). This localization is probably due to the activity of two allelic loci in the diploid genome.

Since the transcription pattern of the cloned *Avi-antiHox5* and the sense RNA probe pattern were very similar, we must have detected the same antisense RNA with the probes to its 3’ and 5’ ends. Fig. 8 D demonstrates the larval expression, as revealed with the sense RNA probe (Fig. 8 D (a-e)) and as compared to *Avi-Hox5* mRNA pattern (Fig. 8 D (f - j)). It can be seen that the territories of antisense and sense RNA transcription overlap. By the late metatrochophore stage (168 hpf) the antisense transcription spreads to the ectoderm of the second segment and is initiated in the mesoderm (dashed line) (Fig. 8 D (d)). By the nectochaete stage the ectodermal expression of antisense RNA gradually vanishes, while the mesodermal transcription in the third segment and future growth zone intensifies (Fig. 8 D (e)). During the same period *Avi-Hox5* mRNA is intensively expressed in the ectoderm of the third chaetiger of the larva but not in the mesoderm or the growth zone (Fig. 8 D (i, j)).

Both antisense transcripts are detected on the territory of the future third larval chaetiger and overlap with the expression pattern of *Avi-Lox5* mRNA (described in Kulakova et al., 2007) starting from the middle trochophore stage. We performed double WMISH with the probes to the cloned *Avi-antiHox5* (Dig-probe) and *Avi-Lox5* (FITC-probe), using BM-purple and Fast Red, respectively, for the signal detection. Based on the results of chromogenic detection, we cannot be sure that the signals are co-localized, but the borders of transcription domains of *Avi-antiHox5* and *Avi-Lox5* coincide (Fig. 9 A).

**Figure 9.**
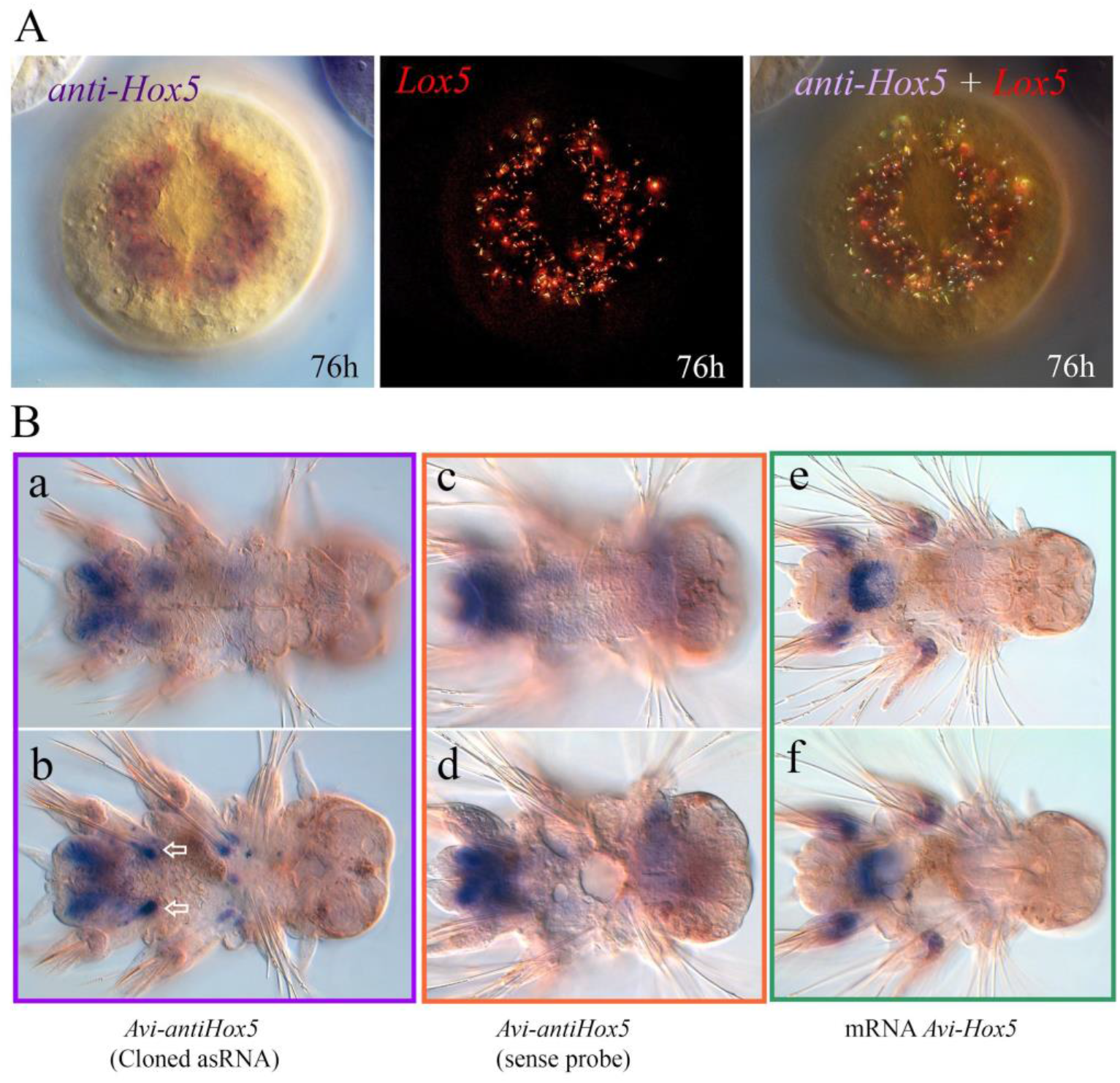
Co-localization of *Avi-antiHox5* and *Avi-Lox5* in *A. virens* trochophore. Sense and antisense transcription of the *Hox5* gene in *A. virens* larvae. **A.** Double WMISH with the probe to *Avi-antiHox5* and antisense strand as probe to *Avi-Lox5* at the stage of middle trochophore (76 h). DIG- and FITC-probes to *Avi-antiHox5* and *Avi-Lox5* are detected with BM-Purple and FastRed, respectively. Vegetal view, ventral side to the top. **B.** WMISH with the probe to *Avi-antiHox5* (purple framework) at the nectochaete stage. White outline arrows (**b**) indicate the unique *Avi-antiHox5* expression at the basis of aciculae (supporting chaetae) (**a**, **b**). WMISH with sense strand as probe detects *Avi-antiHox5* transcript (orange framework) at the nectochaete stage (**c**, **d**); WMISH with antisense strand as probe detects *Avi-Hox5* transcript (green framework) (**e**, **f**); on **B** (**b**), (**d**), (**f**) present deep optical slices. The full description of the transcription pattern is in the text.

The difference between the expression patterns of *Avi-antiHox5* detected with the sense RNA probe and the pattern of the cloned antisense fragment can be seen for the first time at the nectochaete stage (Fig. 9 B). Both antisense transcripts are detected in the growth zone and, less intensively, in the third larval segment. The sense RNA probe signal demonstrates mesodermal localization. Moreover, a weak expression in the pharynx is observed (Fig. 9 B (c, d)). The cloned *Avi-antiHox5* transcript is more distinct in the ectoderm (Fig. 9 B (a)). Besides, its expression is initiated in the basis of aciculae (large supporting rods in parapodia) (Fig. 9 B (b)). This is a transient transcription domain, which is not observed in juvenile worms (Fig. 10). *Avi-Hox5* mRNA expression maximum is observed in the ganglia of the third larval chaetiger, while both antisense transcripts demonstrate a low expression level there (Fig. 9 B (e, f)).

**Figure 10.**
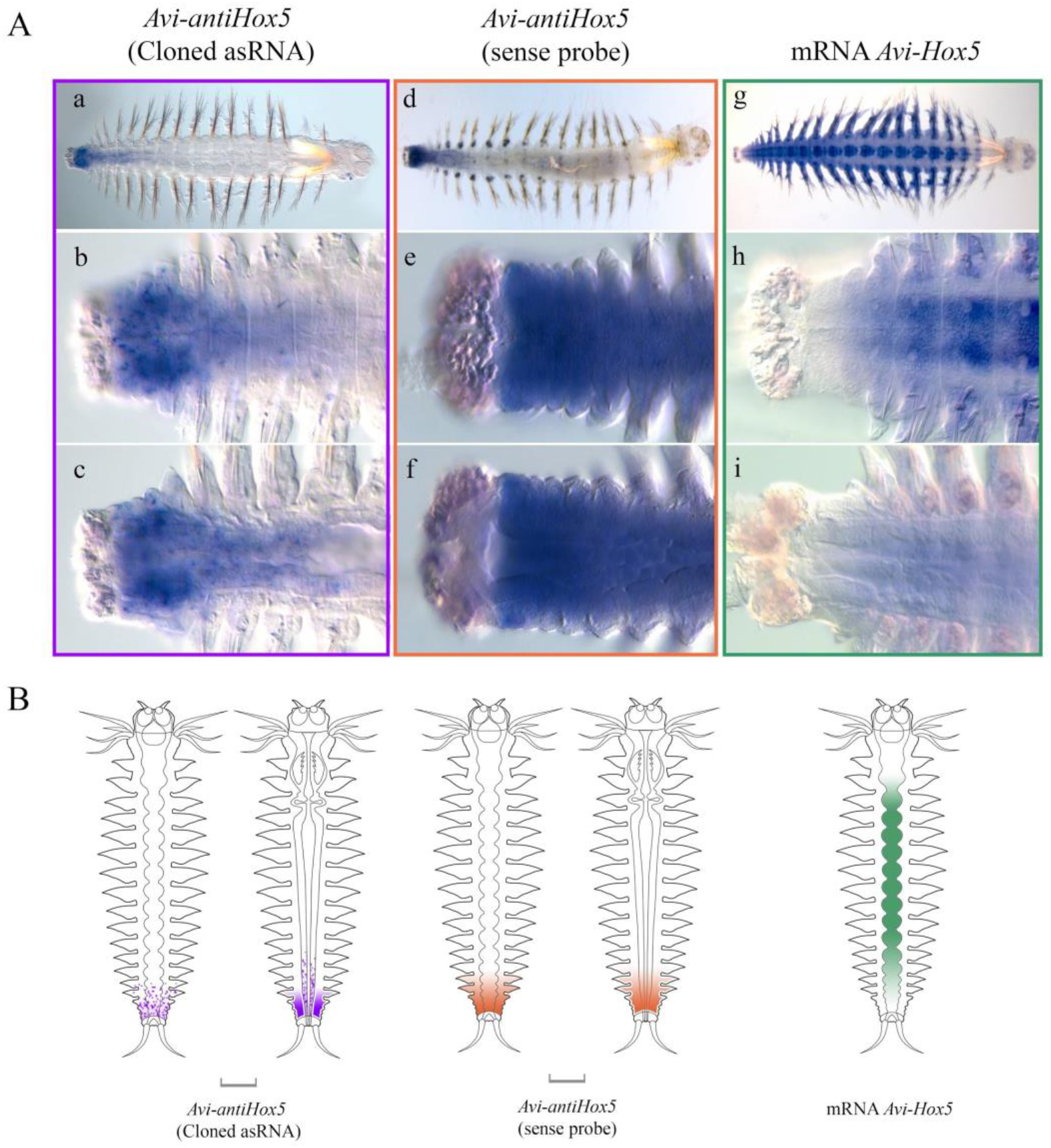
Sense and antisense transcription of the *Hox5* gene in juvenile *A. virens*. **A.** WMISH with the probe to cloned *Avi-antiHox5* (**a**-**c**, purple framework), sense strand as probe to *Avi-Hox5* mRNA (**d**-**f**; orange framework) and antisense strand as probe to the same sequence (**g**-**i**, green frame). Head is to the right on all photos. Pygidium, growth zone and young segments are shown from the ventral side on **b**, **e**, **h** and on the deep optic slices on **c**, **f**, **i**. **B.** Schematic representation of expression patterns of all studied transcripts. The full description of the transcription pattern is in the text.

In juvenile worms both antisense transcripts are revealed in the derivatives of all three germ layers and demonstrate a posterior-anterior expression gradient (Fig. 10 A (a - f)). In the nervous system the gradients are very short and look complementary to the mRNA gradient (Fig. 10 A (h)). The cloned *Avi-antiHox5* fragment is visible in multiple individual cells in the neural system, the surface epithelium, the coelom and the intestine (Fig. 10 A (b, c)). The pattern revealed by sense RNA probe is diffuse and restricted to the ectoderm in a few young segments (Fig. 10 A (e, f)). *Avi-Hox5* mRNA is expressed in a wide bidirectional gradient, with the expression maximum localized in the segments of the central part of the postlarval body (Fig. 10 A (g)) (Bakalenko et al., 2013). *Avi-Hox5* mRNA is detected only in the surface epithelium and in the neural ganglia of the segments that have already formed parapodia. This is clearly visible in optical sections (Fig. 10 A (h, i)). The schematic patterns are presented in Fig. 10 B. Thus, the coding *Avi-Hox5* RNA and its antisense transcription seem to work in overlapping territories at the trochophore and the metatrochophore stages and in adjacent territories in the nectochaete and the juvenile worm.

The *antiHox5* transcription of *P. dumerilii* is detected with the sense RNA probe, which coincides with the coding mRNA at the level of the first and the second exons excluding the activator domain, 5’UTR and 3’UTR (Fig. 11 A). We analyzed the expression of *Pdum-antiHox5* in juvenile worms only and revealed the pattern complementary to the sense one (Fig. 11 B, C). The antisense RNA is mainly detected in the ectodermal growth zone, the surface epithelium and in the neural system of the youngest segments. The diffuse signal is similar to the ectodermal pattern of the non-cloned asRNA of *A. virens* (Fig. 10 A (d, e, f)).

**Figure 11.**
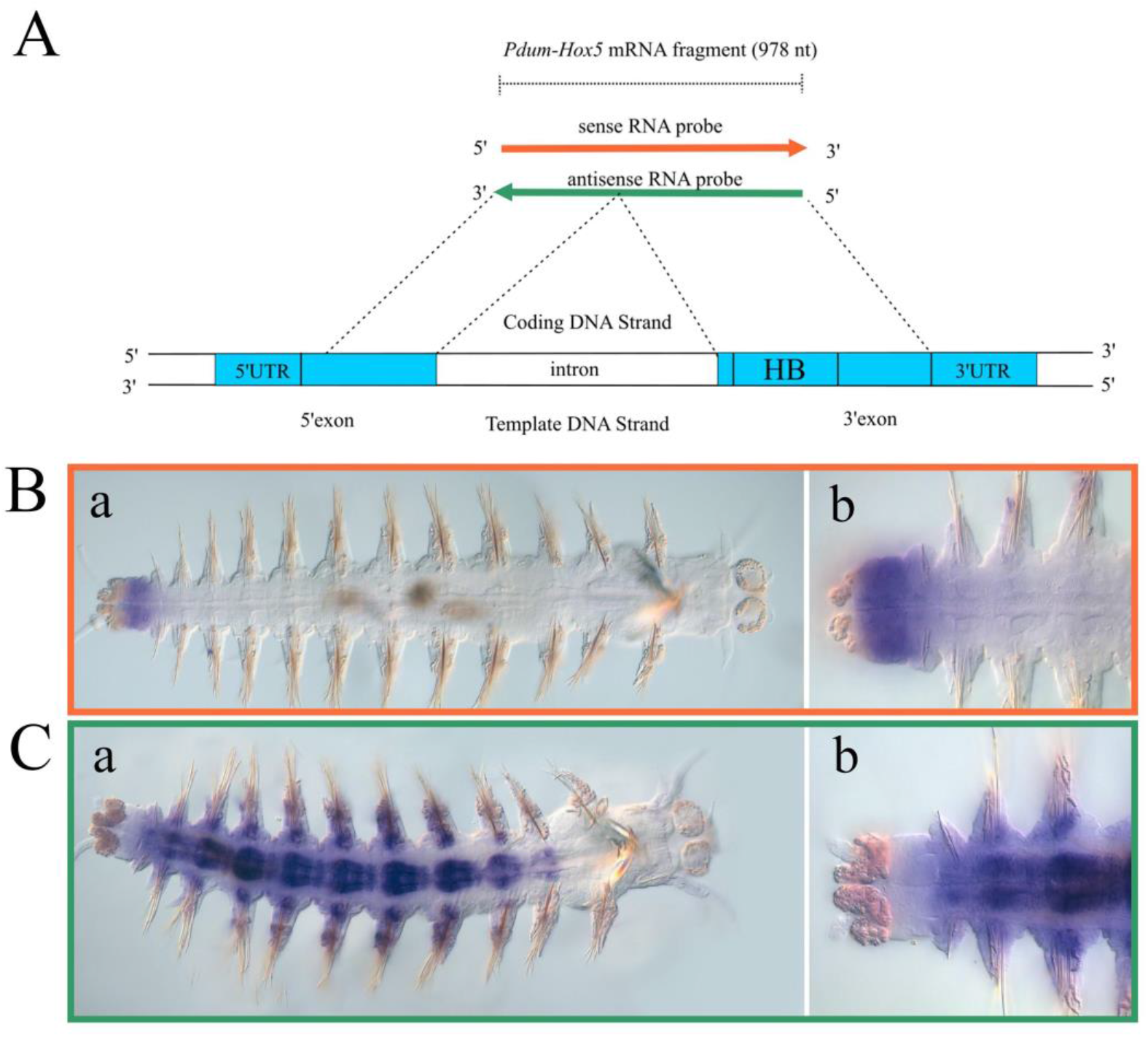
Sense and antisense transcription of the *Hox5* gene in juvenile *P. dumerilii*. **A.** Schematic representation of the probes’ position with the projection to genomic sequence. **B.** WMISH with sense strand as probe to reveal *Pdum-Hox5* antisense transcript (orange framework). **C.** WMISH with antisense strand as probe to reveal *Pdum-Hox5* mRNA (green framework). The full description of the transcription pattern is in the text.

#### antiHox7

We have previously described the antisense transcription of the Hox7 gene of *Alitta virens* during postlarval growth and regeneration (Bakalenko et al., 2013; Novikova et al., 2013). We used the sense RNA probe (522 bp), which coincided with the coding sequence of *Avi-Hox7* mRNA at the level of the second exon (Fig. 12 A). According to our new data, the antisense transcript starts to express much earlier than the sense one. We first detect it at the late trochophore stage (100 hpf) in the vegetal plate cells (Fig. S1.(a)). Later, the transcript appears in the forming stomodeum and in the neural ectoderm of the third segment of the metatrochophore (Fig. S1 (b)). The antisense RNA is detected in the formed ganglia starting from the third larval segment, and its localization does not coincide with the sense transcription area, as *Avi-Hox7* mRNA is observed in the growth zone of nectochaetes and juvenile worms at the same time (Fig. 12 C (a), D (a)). In older worms the sense and the antisense territories overlap in the nervous system (Fig. 12 C (b, c), D (b, c)). The antisense RNA forms a wide anterior-posterior gradient, with the maximum in the anterior third part of the postlarval body. There is no transcription in the growth zone (Fig. 12 C (b, c), B). The mRNA *Avi-Hox7* is revealed in the nervous system and the segment ectoderm in the form of posterior-anterior gradient, the expression in the nervous system vanishing in the anterior third part of the body (Fig. 12 D (b, c), B). On the territory where *Avi-antiHox7* and *Avi-Hox7* patterns overlap, their transcription picture in ganglia is not identical (Fig. 12 C (e), D (e)). Using 5’RACE method, we cloned the fragment of *Avi-antiHox7* lncRNA 1244 nt long, which is localized downstream of 3’-area of *Avi-Hox7* and overlaps with its 3’UTR (Fig. 12 A, purple color). WMISH with the probe to this fragment did not reveal any transcription.

**Figure 12.**
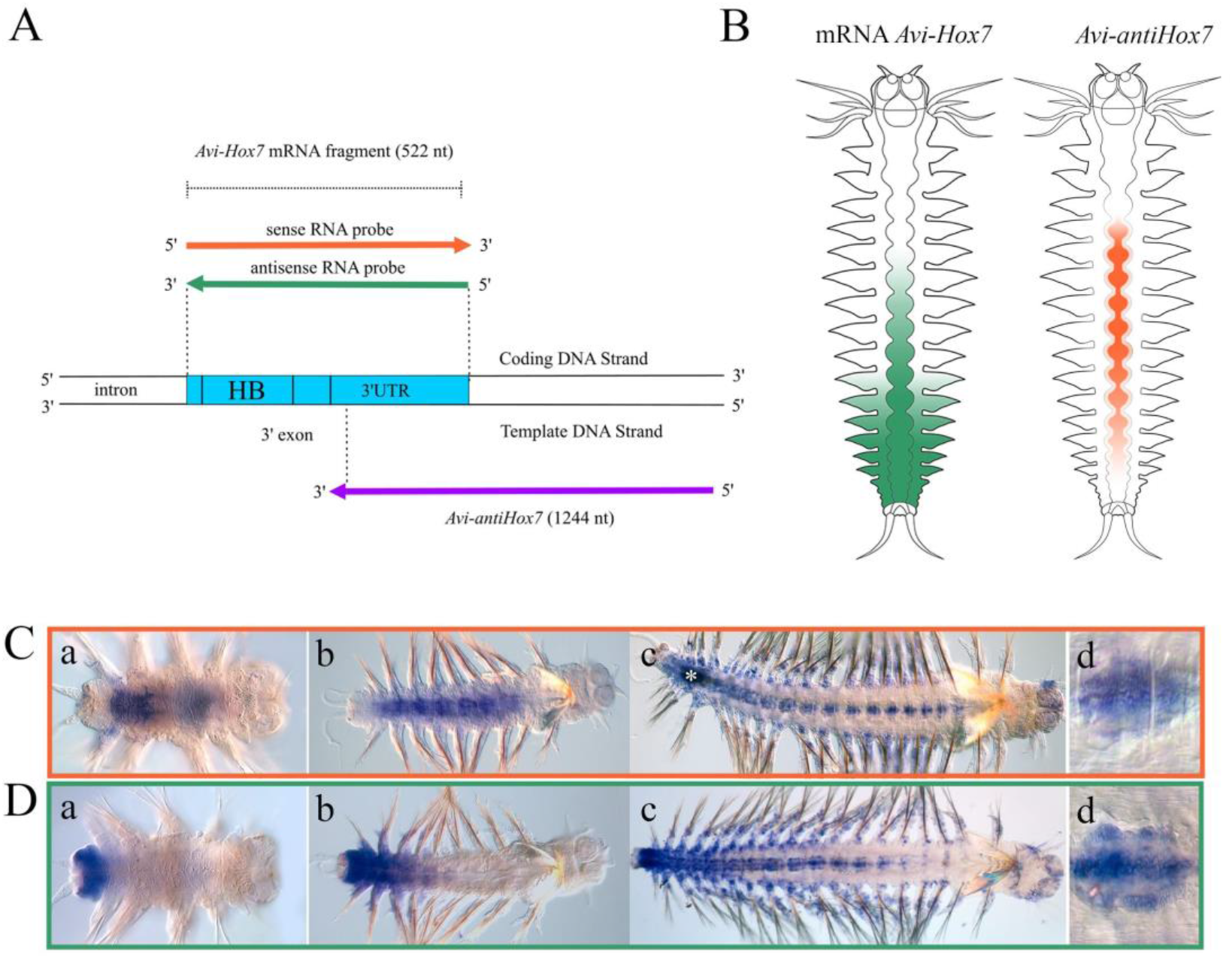
Sense and antisense transcription of the *Hox7* gene in juvenile worms of *A. virens*. **A.** Schematic representation of the probes used with their projection onto the genomic sequence. The purple color indicates the fragment of the cloned *Avi-antiHox7* (5’RACE), which probe does not reveal any transcription. **B.** Schematic expression patterns of studied transcripts of *Avi-Hox7.* **C.** WMISH with the sense strand as probe (**a**-**d**; orange frame). **D.** WMISH with the antisense strand as probe to reveal mRNA of *Avi-Hox7* (a-b; green framework). On **C (d)** and **D (d)** the transcription pattern of antisense and sense RNA in the ventral ganglion of the formed segment (from the middle part of the worm) is shown. The white asterisk on **C (c)** marks the unspecific background in the intestine. The full description of the transcription pattern is in the text.

For *P. dumerilii* we used the sense probe to *Pdum-Hox7* mRNA and the probe to the cloned fragment of *Pdum-antiHox7*. No clear result was identified with any of these probes, perhaps, due to insufficient sensitivity of the method.

#### antiLox4

We did not detect the antisense transcription of *Avi-Lox4* gene using the sense RNA probe. The size of the probe we used (302 nt) and/or the area of its overlapping with the antisense transcript are probably too small to detect the transcription with WMISH (Table S2).

However, we found the antisense transcription of *Pdum-Lox4.* Sense RNA probe to 3’-end of mRNA corresponds to a homeobox fragment and 3’-UTR (Fig. 13 A). It reveals *antiLox4* transcription in coeloms of several last segments (Fig. 13 C (a, b), B). The probe to 3’ - fragments of the sense RNA detect the expression in neural ganglia and segmental ectoderm in the form of a wide posterior-anterior gradient (Fig. 13 D (a, b), B).

**Figure 13.**
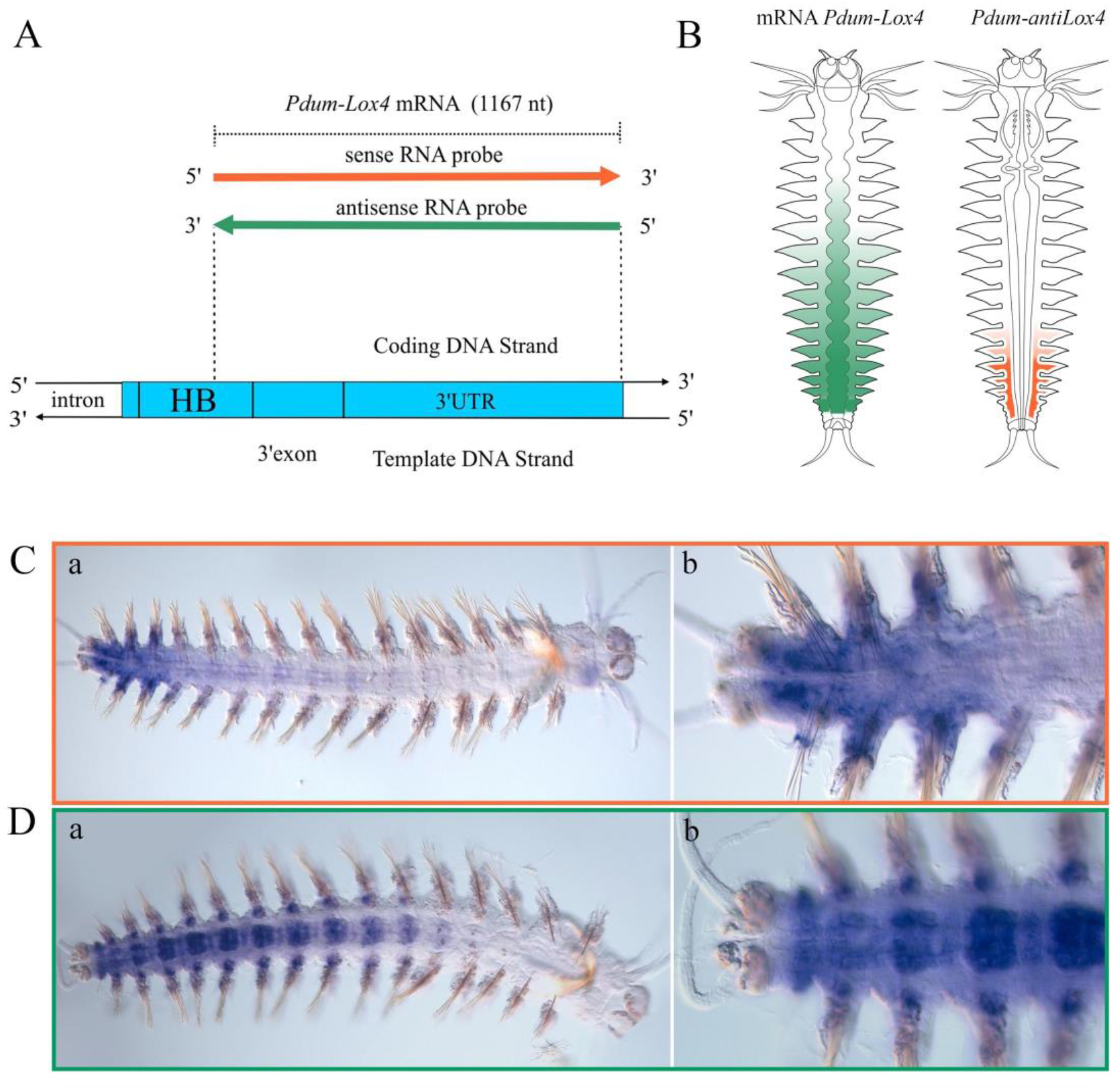
Sense and antisense transcription of the *Lox4* gene in juvenile *P. dumerilii*. **A.** Schematic representation of the probes used to 5’ and 3’-areas of mRNA with their projection onto the genomic sequence. **B.** Schematic representation of expression patterns of studied *Pdum-Lox4* transcripts. **C**. WMISH with sense strand as probe to 3’-area of *Pdum-antiLox4* transcript (**a**, **b**; orange framework). **D.** WMISH with antisense strand as probe to detect *Pdum-Lox4* transcript (**a**, **b**; green framework). The full description of the transcription pattern is in the text.

#### antiLox2

As in the previous case, we did not reveal the antisense transcription of *A. virens Lox2* with the sense RNA probe (516 nt) that corresponds to the 3’-end of the protein-coding sequence and 3’-UTR (Table S2). However, there is an antisense transcript of *P. dumerilii Lox2* that was detected with the sense RNA probe (720 nt). This probe coincides with 5’UTR sequence and the coding part of the first and partially second exons (Fig. 14 A). Antisense transcription is detected in the caudal part of the intestine and adjacent coelomic epithelium in the youngest segments (Fig. 14 B, C (a, b);). *Pdum-Lox2* mRNA is expressed as a gradient in neural ganglia and in the ectoderm of young segments (Fig. 14 B, D (a, b)). Interestingly, at the early stages of regeneration (10 hours post tail amputation) *Pdum-antiLox2* starts to express in the neural ganglion close to the amputation site (Fig. 14 C (c, d)). This can be explained either by the changes in the function of the antisense transcript during the regeneration process or, more likely, by an intensification of the neural system expression, which makes it possible to detect the domain invisible during the normal growth.

**Figure 14.**
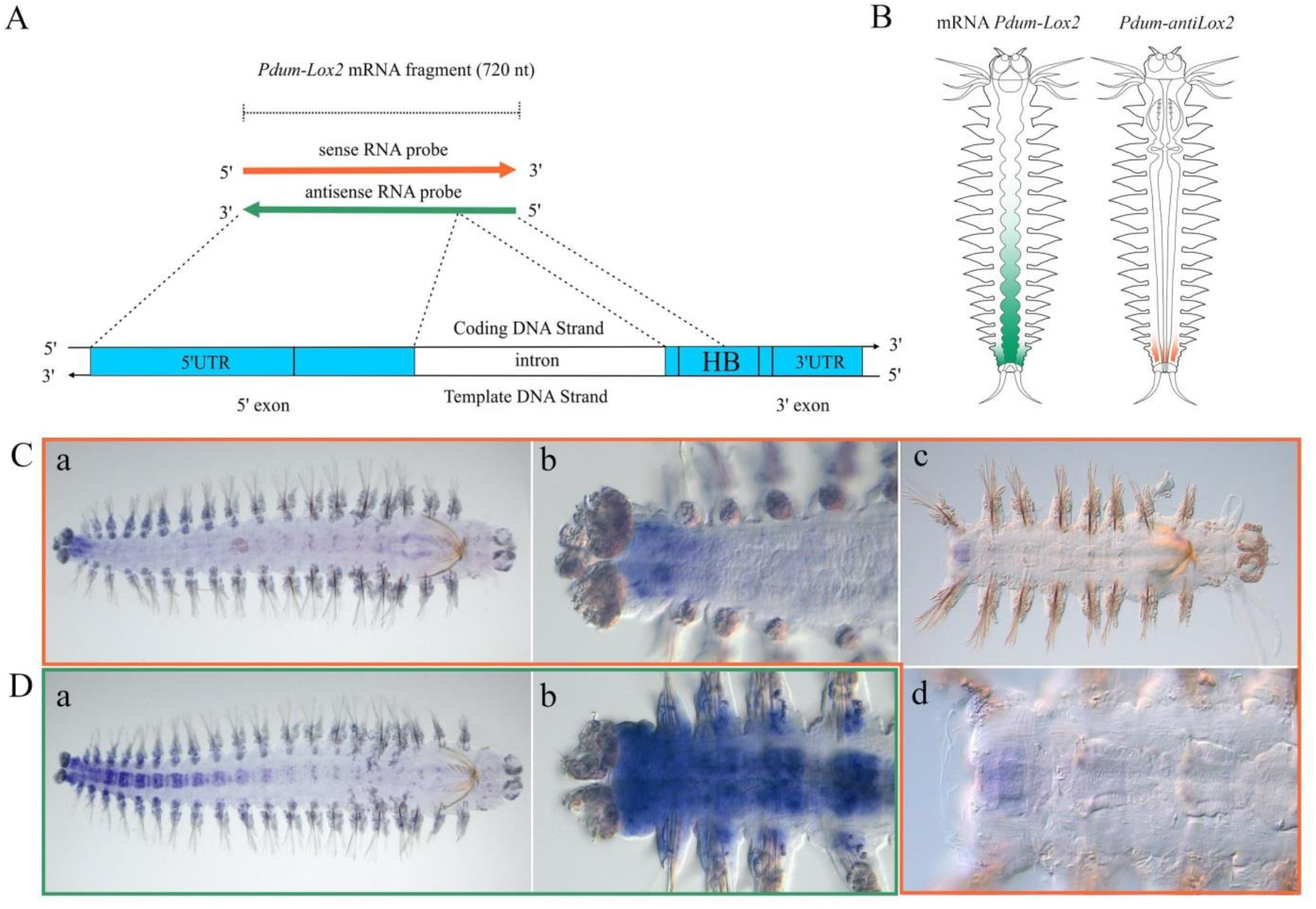
Sense and antisense transcription of the *Lox2* gene in juvenile *P. dumerilii*. **A.** Schematic representation of the probes used with their projection onto the genomic sequence. **B.** Schematic representation of expression patterns of studied *Pdum-Lox2* transcripts. **C.** WMISH with the sense strand as probe to reveal *Pdum-antiLox2* transcription (**a** - **d**; orange frame) **D.** WMISH with the antisense strand as probe to reveal *Pdum-Lox2* transcription (**a**, **b**; green framework). On **C (c)** and (**d)** the expression of *Pdum-antiLox2* in the ganglion of the regenerating worm (10 hours post amputation) is shown. The full description of the transcription pattern is in the text.

#### antiPost2

To detect the antisense transcription of *Avi-Post2* gene (GenBank: KY020041.1) we used sense RNA probe (706 nt), which coincides with the first exon and a small part of the second one (Fig. 15 A). *Avi-antiPost2* can be revealed only on larval stages during a short period starting from the middle trochophore stage (70 hpf) and to the early metatrochophore stage (112 hpf) (Fig. 15 B (a-d)). This transcript is visible presumably in the nuclei of the large cells localized on the vegetal pole of the larva (Fig. 15 C (a, b)). Later the descendants of these cells will become a part of the pygidium.

**Figure 15.**
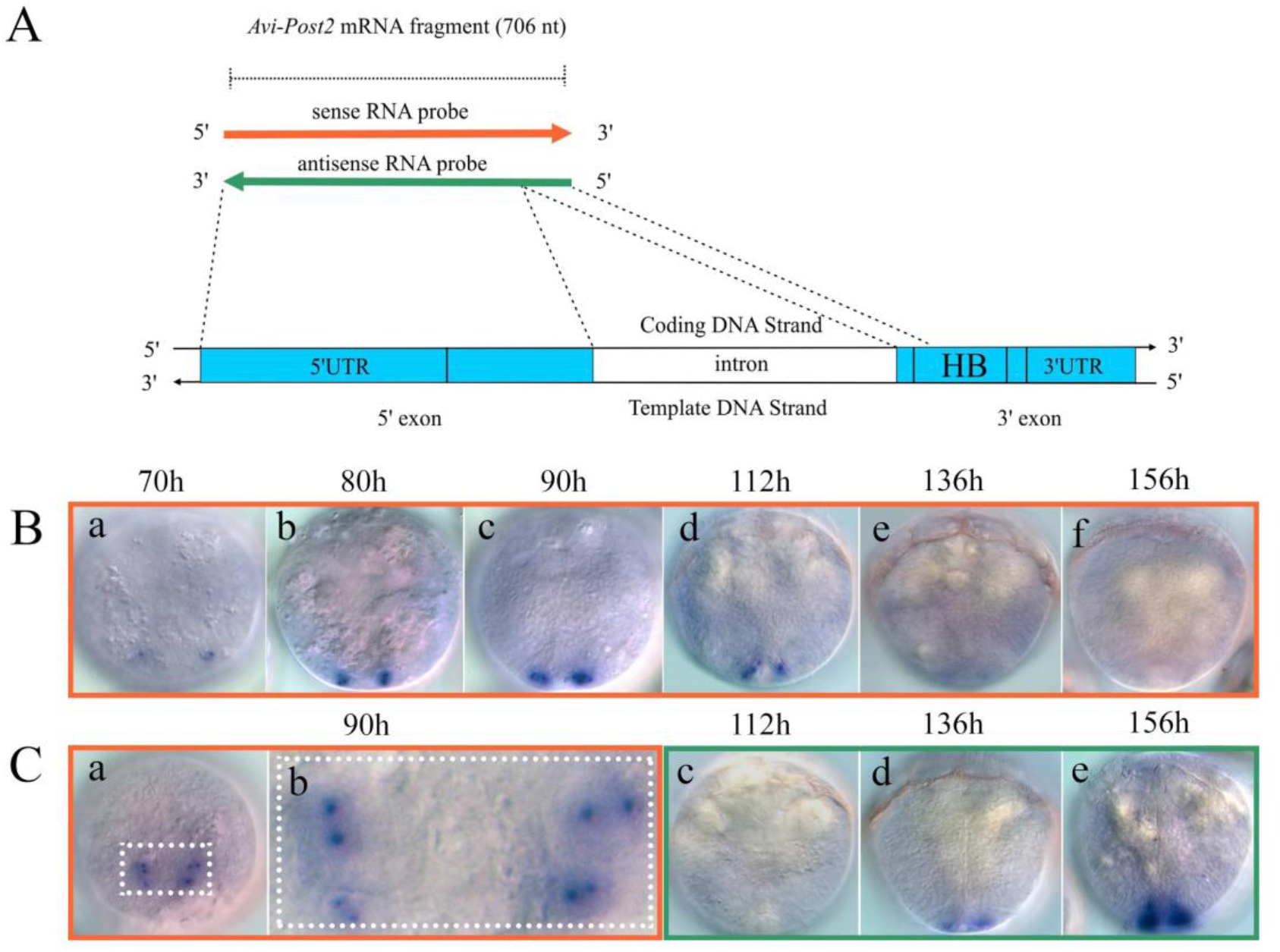
Sense and antisense transcription of the *Post2* gene in *A. virens* larvae. **A.** Schematic representation of the probes used with their projection onto the genomic sequence. **B.** WMISH with the sense strand as probe to detect *Avi-antiPost2* transcription (**a**-**f**, orange framework). **C** (**a**-**b**). WMISH with the sense strand as probe to detect *Avi-antiPost2* transcription (orange framework). **C** (**c**-**e**). WMISH with the antisense strand as probe to detect *Avi-Post2* transcription (green framework). All larvae have frontal orientation. **C** (**a**) presents the 90 hpf larva, view from the vegetal pole. **C** (**b**) (dotted frame) demonstrates the enlarged fragment of **C** (**a**) with the signal in two dots presumably in the nuclear. The full description of the transcription pattern is in the text.

At the late trochophore stage the transcript spreads all over the cell, vanishing completely by the middle metatrochophore stage (Fig. 15 B (e, f)). At this time point *Avi-Post2* mRNA is synthesized in the region of the future pygidium (Fig. 15 C (d, e)). It is difficult to say if the cells that synthesized the antisense transcript before switch to the synthesis of the sense transcript, since there is a pause between the vanishing of one transcript and the activation of the other. To answer this question, we need to increase the sensitivity of the method and to use more frequent fixations.

The antisense transcript of *Pdum-Post2* is detected with the sense RNA probe (750 nt), that almost completely coincides with the second exon of this gene (Fig. 16 A). Similarly to *Avi-antiPost2*, this antisense RNA starts to be synthesized presumably in the nuclei on the territory of the future pygidium long before the activation of the sense transcript. At the early metatrochophore stage, when *Pdum-Post2* mRNA expression is initiated in pygidial lobes, the antisense transcript is still detected. It is localized in the adjacent area at the level of the trochoblasts of the telotroch, i.e. around the future pygidial lobes (Fig. 16 B (a), C (a)). This transcription gradually vanishes, but by the stage of late metatrochophore/early nectochaete the new *Pdum-antiPost2* expression domain appears in the forming pharynx (Fig. 16 B (b)). *Pdum-Post2* mRNA transcription retains in the pygidium (Fig. 16 C (b)). At this stage the sense and antisense transcripts are synthesized not in the adjacent territories but in separate ones, similarly to what we observed in case of sense and antisense *Pdum-Hox3* RNA (Fig. 4 B). This may point to the independent function of *Pdum-antiPost2* in the development, which is not connected with the control of the transcription and/or translation of the sense RNA. Juvenile worms do not transcribe *Pdum-antiPost2* at all (or not on the level detectable by the method we use). However, during regeneration in 48 hours after tail amputation we observe the antisense transcription in the ectoderm of the pygidium anlagen again (Fig. 16 B (c, d)). The sense transcript is detected at this stage mainly in the inner cell mass of regeneration blastema and overlaps a little with the antisense transcription (Fig. 16 C (c, d)).

**Figure 16.**
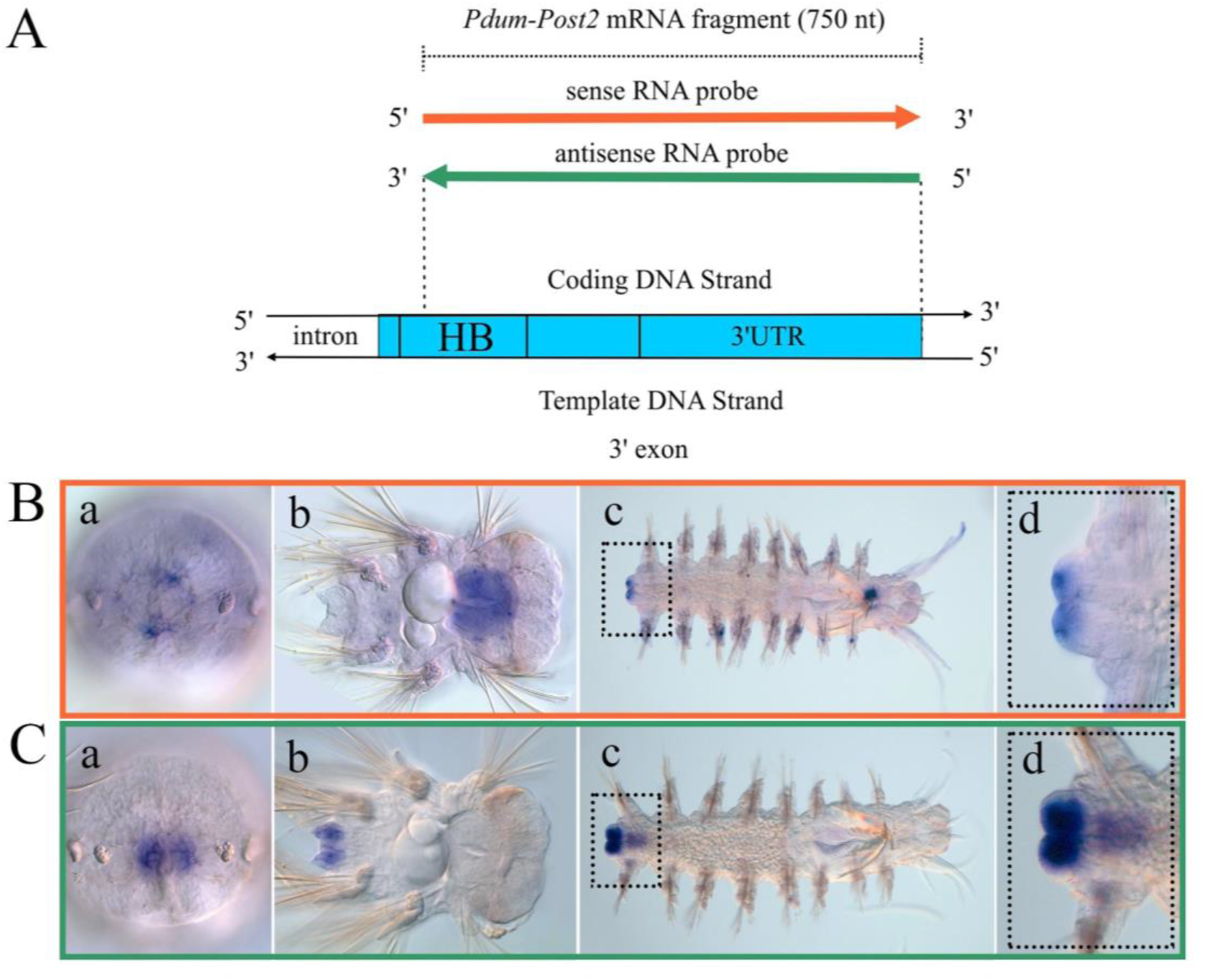
Sense and antisense transcription of the *Post2* gene in larvae and juvenile worms of *P. dumerilii*. **A.** Schematic representation of the probes used with their projection onto the genomic sequence. **B.** WMISH with the sense strand as probe to detect *Pdum-antiPost2* (**a**-**d**; orange frame). **C.** WMISH with the antisense strand as probe to detect *Pdum-Post2* (**a**-**d**; green frame). On **B** (**a**) and **C** (**a**) the metatrochophores from the vegetal pole are shown. Dotted frames on **B** (**c)** and (**d)** and **C (c)** and (**d)** indicate the antisense and sense transcription in the regenerating worm (2 days post amputation), respectively. The full description of the transcription pattern is in the text.

## Discussion

### 1. Almost all Hox genes of nereidid annelids possess antisense transcripts

We investigated the specific transcription patterns of Hox genes’ NATs of nereidid annelids by WMISH using either probes to the cloned antisense RNAs or sense RNA probes to mRNAs. Figure 17 (Fig. 17) summarizes what kinds of probes were used to analyze transcription of different Hox gene RNAs and indicates the sequences used for probe synthesis. Our research shows that despite the difficulties of working with lncRNAs due to their low copy number and a short lifespan, there is a way to study their transcription patterns even without the cloning of the gene fragment. If NATs overlap the coding sequence for more than 500 nt, it is possible to use sense RNA probes for the analysis of the genes of interest (Table S2).

**Figure 17.**
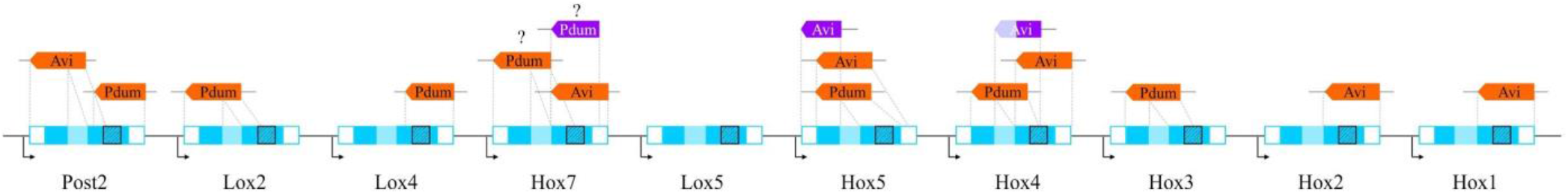
Schematic representation of presumptive Pdum and Avi Hox cluster with the projections of probes’ positions. Orange color indicates sense probes to mRNAs of corresponding Hox genes. Lavender color indicates the probes to the cloned NATs of Hox genes. The scheme contains the probes that detected expression patterns and two probes (marked by question mark) that gave controversial results for which more analysis is regarded. Blue outline marks the sequences of Hox genes. Shading: white – 5’ and 3’-UTRs; blue – protein coding areas; light blue – introns; shaded boxes – homeobox areas.

We used probes to 5’- and 3’-areas of Hox genes and revealed asRNAs for the genes of all paralogous groups except *Lox5* (Table S2). We cannot say with certainty that this Hox gene does not possess asRNA, but the sense RNA probes to the 5’area of *Pdum-Lox5* and the 3’-area of *Avi-Lox5* did not reveal any expression. All asRNAs of Hox genes, whose patterns were described in this work, can be referred to as NATs (Ariel et al., 2015).

The exons of cloned NATs *(Avi-antiHox4_1, Avi-antiHox4_2, Avi-antiHox7)* consist not only of the coding sequences of Hox genes but also of the intronic and even intergenic regions (Fig. 1). NATs of mammalian Hox genes are organized in a similar way (Mainguy et al., 2007). Besides, we cannot rule out the presence of polycistronic transcripts whose exons overlap with several Hox sequences simultaneously. Such polycistronic sense and antisense RNAs are described for human and murine Hox clusters (Mainguy et al., 2007). In case of in situ screening, RNAs like that should be revealed by the probes to different Hox transcripts but should demonstrate similar expression patterns. We show that all the described patterns are gene-specific, however, asRNAs of several different Hox genes are detected in the same areas, the esophagus and the stomodeum (Fig. 2 B, C (a); Fig.3 B, C (c); Fig.7 B (b); Fig. 16 B (b)).

The probes to the cloned antisense sequences and sense RNA probes to the same gene demonstrated variable expression patterns (Fig. 5, 6 *(Avi-Hox4);* Fig. 9, B *(Avi-Hox5)).* This means that there are at least three different antisense transcripts for *Avi-Hox4* (including *Avi-Hox4_2*) and two for *Avi-Hox5*. It is still an open question, how many different asRNAs are transcribed from Hox clusters of nereidids. To answer it, the full sequencing of the clusters and the cap analysis of gene expression (CAGE) are needed.

### 2. NATs’ patterns of nereidid Hox genes are gene-specific and closely related to sense transcription patterns

NATs’ patterns of *A. virens* and *P. dumerilii* Hox genes are so variable and dynamic that at first sight it seems impossible to elucidate the common principles of their behavior. To introduce some order into the confusing abundance of data, we decided to consider NATs as possible effectors influencing the functioning of their complementary mRNAs. In this case, we can suggest five possible variants of sense/antisense interaction.

#### a) Complementary transcription on adjacent territories

This option is realized for the majority of sense Hox RNAs and their NATs. In juvenile worms and nectochaetes (if this stage was studied) the expression patterns of *Avi-antiHox1*, *Avi-antiHox2, Avi/Pdum-antiHox4, Avi/Pdum-antiHox5*, *Pdum-antiLox4* и *Pdum-antiLox2* are complementary to the sense patterns. This means that the adjacent territories of two transcripts (sense and antisense) do not overlap or overlap on small territories. *Avi-Hox2* and *Avi-antiHox2* are an example of non-overlapping transcription domains. Both transcripts are detected in the growth zone and in the anterior parts of the digestive tract, but sense RNA is localized in the mesodermal component of the growth zone and at the basis of the jaws, while its NAT is located in the ectodermal growth zone and the esophagus (Fig. 3 C, D). *Avi/Pdum-Hox5* and *Avi/Pdum-antiHox5* transcripts are distributed in the form of opposite gradients and have a narrow conjoint area in the segmental ectoderm and the neural system (Fig. 10, 11).

Noteworthy, *Avi-antiHox7* is expressed according to this rule at the nectochaete stage and in juvenile worms (4-5 segments), but as the animal grows the areas of sense and antisense expression start to overlap considerably in the neural system due to the activation of sense RNA in postlarval segments (Fig. 12 B, C, D). The expression is probably still complementary but on the level of individual cells, since the pattern in the ganglia differs for the sense and the asRNAs (Fig. 12 C (d), D (d)) (Bakalenko et al., 2013). *Pdum-Post2/Pdum-antiPost2* transcripts display complementary transcription only at the stage of the metatrochophore (Fig. 16 B (a), C (a)). Later the complementary transcription is observed only during regeneration (Fig. 16 B (c, d), C (c, d)). Noteworthy, the patterns of different asRNAs of *Avi-Hox4* are complementary not only to the sense transcript but also to each other (Fig. 6 A, B).

#### b) Early antisense transcription that precedes and potentially prevents the sense one

Transcription of NATs of *Avi-Hox5*, *Avi-Hox7* and *Avi/Pdum-Post2* is initiated at the trochophore stage, long before the activation of their sense RNAs. In case of *Avi-Hox5* both antisense transcripts precede the expression of mRNA in the third larval segment. Later all the three transcripts are co-expressed in this territory until the nectochaete stage (Fig. 8). The early transcription of *Avi-antiPost2* precedes the expression of sense RNA at the same domain and on the adjacent territory, but antisense transcription vanishes before mRNA starts to express (Fig. 15 B, C (c-e)). *Pdum-antiPost2* demonstrates a very similar early pattern, which is retained after the initiation of sense transcription, and two expression areas seem to be strictly complementary at the stage of early metatrochophore (Fig. 16 B (a), C (a)). *Avi-antiHox7* transcript appears in larval neuroectoderm at the level of the third segment (Fig. 12 C (a), Fig. S1). Sense transcription never spreads to the territory of the third larval segment, neither at later stages nor even during regeneration, when the anterior boundary of *Avi-Hox7* expression domain spreads anteriorly (Novikova et al., 2013).

#### c) Overlap of sense and antisense RNA patterns

mRNAs and their NATs of nereidid Hox genes can be temporarily or permanently transcribed at the same territory. The first variant is typical for the larval transcription of *Avi/Pdum-antiHox4* and *Avi-antiHox5* (Fig. 5 B (b), C (b); Fig. 7 B (a), C (a); Fig. 8 D (d), E (d)). Sense and antisense RNAs of *Avi-Hox1* permanently overlap in the esophagus (Fig. 2 B, C (a), D (a)).

#### d) Transcription in spatially divided regions

In some cases we observe an independent transcription of mRNAs and their NATs on non-adjacent territories: *Pdum-Hox3* (growth zone) and *Pdum-antiHox3* (esophagus) (Fig. 4 B); *Pdum-Hox4* (ectoderm of the second and the third nectochaete segment) and *Pdum-antiHox4* (stomodeum and distinct cells in the episphere) (Fig. 7 B (b), C (b)); *Pdum-Post2* (pygidium) and *Pdum-antiPost2* (stomodeum) (Fig. 16 B (b), C (b)). We suggest that in this way NATs prevent a non-specific expression of their mRNAs. Conversely, these observations may indicate an independent function in the regulation of alternative targets, which is usually the case for trans-NATs (Deforges et al., 2019).

#### e) Transcription on the territory of the neighboring Hox gene expression domains

The expression patterns of some NATs coincide with the sense patterns of the neighboring Hox genes. For example, *Avi-antiHox2* is expressed in the ring of ectodermal cells where *Avi-Hox3* works (Fig. 3 B (a, b), C (a, b); Fig.4 B (b)) (Bakalenko et al., 2013). At one of the larval stages *Avi-antiHox5* and *Avi-Lox5* are localized at the same territory of the future third larval segment, which was confirmed by double WMISH (Fig. 9 A). A pattern like this may indicate the presence of bidirectional transcription, which is initiated from the closely localized promoters on the opposite DNA strands. These promoters are under the common regulation, thus the transcribed RNAs are co-expressed (Balbin et al., 2015). A detailed analysis of this observation is impossible without the data on the structure of nereidid Hox cluster. It is known that *A. virens* and *P. dumerilii* Hox genes are located in one locus but the order of the genes and the presence of inversions in the clusters are unknown (Andreeva et al., 2001; Hui et al., 2012).

### 3. Putative functions of nereidid Hox genes’ NATs and their possible role in development

LncRNAs are encoded in the Hox clusters of of mammals, hemichordates, insects and myriapods (Casaca et al., 2018; Mainguy et al., 2007; Freeman et al., 2012; Pettini and Ronshaugen, 2016; Petruk et al., 2006; Brena et al., 2006). These molecules not only control the transcription of Hox genes but also can be involved in the other important processes such as the switch of cellular metabolism from glycolysis to oxidative phosphorylation (Huang et al., 2017). The best studied among them are long intergenic noncoding (linc)RNAs, for example HOTAIR and HOTTIP in mammals (Gupta et al, 2010; Balbin et al., 2015; Li et al., 2019).

LncRNAs of nereidid Hox genes belong to another class of molecules. These are NATs (Natural Antisense Transcripts), and they are much more poorly studied in model animals. Human Hox clusters are known to code dozens of NATs (Mainguy et al., 2007), some of which are conservative at least between human and mouse, and so cannot be considered as transcriptional noise (Mainguy et al., 2007; database LncBook: https://bigd.big.ac.cn/lncbook/index). These transcripts overlap Hox genes’ sequences on the regions from 44 nt to 607 nt, and their functions are not yet properly studied. The transcription of some of these ncRNAs is known to be significantly increased or reduced in tumor cells where they co-express with their target mRNAs and stabilize them from degradation, thus supporting the activity of the target gene (Zhang et al., 2018). We suggest that some NATs of nereidids’ Hox genes can act using a similar mechanism, since their transcription patterns overlap with the expression domains of the mRNAs and since both transcripts are detected in the cytoplasm.

Despite the lack of knowledge concerning the function of antisense transcripts in the Hox cluster, enough data regarding the role of NATs in different regulatory systems have been accumulated. One of the described mechanisms of sense and antisense transcripts’ interplay is transcriptional interference. This mechanism comes into force if sense and antisense transcriptional start points are localized in close proximity to each other. In this case the transcriptional machinery of the antisense strand prevents the synthesis of mRNA by mechanically blocking the polymerase II complex of the sense strand (Latos et al., 2012; Latge et al., 2018; Flippot et al., 2019). Transcriptional interference takes place in cell nuclei. The resulting antisense transcript per se plays no role and eventually degrades. A large number of NATs of nereidids’ Hox genes is indeed detected in the nuclei. This early transcription, which is always larval, usually excludes the synthesis of complementary mRNAs on the same territory (as it was demonstrated for *Avi-antiHox5* and *Avi/Pdum-antiPost2*). However, in case of *Avi-Hox4/Avi-antiHox4_1* the nuclear asRNA signal accompanies the cytoplasmic signal of mRNA (Fig. 5 D (d), E (d)).

The nuclear localization of NATs suggests that there may be other probable mechanisms of transcriptional control. On the one hand, lncRNA molecules may decoy the basic and enhancer-specific transcriptional factors, thus preventing the work of promoters of sense transcripts (Morriss and Cooper, 2017; Flippot et al., 2019). On the other hand, lncRNAs may attract those factors to the promoter and act as guide lncRNAs (Chiu et al., 2018; Flippot et al., 2019). Besides, lncRNAs can function in the nucleus as scaffold molecules (Wang and Chang, 2011). In this case, they recruit to the target site (in cis- or trans-position) not only the individual proteins, but the whole protein complexes of chromatin remodeling, thus changing the chromatin state. Here, as in the previous case, both positive and negative regulation models are possible. It is not implausible that an early (preceding or preventing) expression of NATs of *A. virens* and *P. dumerilii* is realized through one of those mechanisms.

Finally, NATs can be processed to siRNA (short interfering RNAs). NAT-siRNAs were found in mammals, Drosophila, *C. elegans* and yeasts (Zhang et al., 2013; Holoch and Moazed, 2015). The mechanism implies the formation of duplexes between lncRNA and the complementary target RNA with further degradation of long dsRNA precursors to the set of small 21-nt molecules by Dicer (Dicer2 in *D. melanogaster*) ribonuclease. Afterwards the short double-stranded precursors are loaded into RISC-complex and participate in post-transcriptional gene silencing (PTGS). Noteworthy, both the sense and the antisense transcript that form dsRNA are synthesized simultaneously and are processed in the cytoplasm. PTGS in different animals including Drosophila and mammals is closely related to transcriptional gene silencing (TGS) (Pal-Bhadra et al., 2004; Weinberg et al., 2006; Hawkins et al., 2009). It was demonstrated in human cell culture that the antisense small RNA, which is complementary to the promoter region of the target gene, participates in TGS initiation. It is important to note that TGS can be inhibited through the inhibition of RNA polymerase II, which can be a sign of the interaction between siRNA and 5’-region of sense transcript (Weinberg et al., 2006). The mechanisms of PTGS realization in Drosophila and mammals may be at work in annelids as well since in many cases sense and antisense RNAs of nereidids’ Hox genes have been detected in the cytoplasm of the same cells (*Avi-Hox4, Avi/Pdum-Hox5*). It is still debatable whether our models involves TGS by siRNAs.

It is known that transcriptional boundaries of most Hox genes in the postlarval body of nereidids are constantly moving because the worms grow for the most part of their lives. The patterns of many sense/antisense Hox transcripts of *A. virens* and *P. dumerilii* retain a dynamic balance. It is important that though the patterns of two transcripts are complementary, the locus that is the source of both RNAs (sense and antisense) stays active. This means that the down-regulation of sRNA transcription does not occur through TGS.

It is known that Hox activation during regeneration in nereidids takes place approximately in two hours after the injury. This is relatively fast in comparison with amphibians and even planarians (Gardiner et al., 1995; Orii et al., 1999). It is possible that in the postlarval nereidid body the Dicer and Ago “team” functions in the nuclei as it was recently shown for other bilaterian animals (Cernilogar et al., 2011; White et al., 2014; Shuaib et al., 2019). In case the complementary asRNAs are synthesized, these proteins cut Hox gene mRNAs as soon as the latter have been synthesized. Thus, Hox mRNA transcription may be constantly retained but its presence may be controlled by up- and down-regulation of asRNA. This hypothesis clearly explains the fast activation of Hox transcription, but needs to be tested experimentally.

It cannot be ruled out that asRNAs of some Hox genes of nereidids serve for maintaining Hox loci in the active state and for preventing heterochromatization. This allows a fast initiation of the sense transcription if the body is damaged. However, if asRNAs serve to maintain an active locus, it is unclear why they should be transported to the cytoplasm.

It is worth noting that our probes, regarding their position relative to 3’ or 5’-ends of the target transcript, contain a full or partial sequence of the homeobox. In the case when we managed to clone the fragments of asRNA (Table S1), all of them except for *Avi-antiHox5* contained a full or a partial sequence that was complementary to homeobox. This may well be the case for *Avi-antiHox5* too, since it was cloned by the 3’RACE method and its 5’-area is unknown. This means that many NATs of nereidids’ Hox genes overlap the most conservative parts of Hox paralogs and potentially may work in trans-mode.

All NATs of nereidids’ Hox genes demonstrate complicated transcriptional dynamics. Most of them change their localization from the nuclear to the cytoplasmic in the course of development, but not vice versa. We often observed the change of the interaction character in the sense/antisense pairs. The patterns may overlap at the early developmental stages, becoming complementary later on *(Avi-antiHox4_1, Avi-antiHox4* (sense probe) and *Avi-antiHox5)* and vice versa *(Avi-antiHox7).* These reversals can indicate the change of NATs’ functions during the development and/or the change of the mechanisms of functional realization.

The transcription of Hox genes’ mRNAs in nereidids and an unrelated annelid *Capitella* sp. (Kulakova et al., 2007; Fröbius et al., 2008; Bakalenko et al., 2013) displays variable patterns during larval and postlarval development. We suggest that this difference, as well as the restricted regeneration capacity of larval segments, is due to the difference in the epigenetic tuning of Hox loci in larval and postlarval annelid body. We hypothesize that this tuning may be controlled by NATs.

### 4. The occurrence of NATs of Hox genes among Bilateria: an individual specificity of regulation or an ancestral feature?

The first NATs of Hox genes were found and partially cloned in mouse (Hsieh-Li et al., 1995). It was shown that at least four different antisense splice forms are transcribed from *Hoxa11* gene and their transcription patterns are complementary to the patterns of the sense transcripts (Hsieh-Li et al., 1995).

In the lineage of protostomian animals, NATs associated with Hox cluster (antisense transcripts of *Ubx* gene (*aUbx*) were found in three species of myriapods, but not in onychophorans (Brena et al., 2006; Janssen and Budd, 2010; Janssen et al., 2014). Thus, the presence of antisense RNAs was considered as a synapomorphy of Myriapoda (Janssen and Budd, 2010). It should be noted, however, that the structure of onychophoran Hox cluster is yet undescribed, while the cluster of the centipede *Strigamia maritima* is intact and well-ordered (Chipman et al., 2014). Besides, the Hox cluster of *S. maritima* is associated with the gene evx/evenskipped similar to the intact Hox clusters of Deuterostomia (Chipman et al., 2014), which indicates a similarity to the ancestral cluster of arthropods. The expression patterns of *aUbx* are studied in detail in the centipede *S. maritima* and the millipede *Glomeris marginata* (Brena et al., 2006; Janssen and Budd, 2010). In general, the principles of transcription of myriapod aUbx are similar to what we observe for some antisense transcripts in nereidids. The *Ubx* gene is specific for Panarthropoda lineage and its regulation by *aUbx* transcript is probably limited by the myriapod lineage. However, considering the data on vertebrates (Hsieh-Li et al., 1995; Mainguy et al., 2007) and annelids, we can assume that the principle of NATs’ functioning is not the invention of myriapods but something that was inherited from the regulatory system of the common ancestor of Protostomia and Deuterostomia. Hox gene mRNAs transcription is relatively conservative in *A. virens* and *P. dumerilii* (Kulakova et al., 2007), but the expression patterns of NATs of orthologous Hox genes of nereidids demonstrate only a remote resemblance (*Avi-antiPost2* and *Pdum-antiPost*; *Avi-antiHox4* and *Pdum-antiHox4; Avi-antiHox5* and *Pdum-antiHox5).* Since there is a probability that our probes reveal non-homologous asRNAs, we should assert the evolutionary lability of NATs with a certain caution. In general, most of lncRNAs are much less conservative than mRNAs or miRNAs (Jarroux et al., 2017). NATs transcribed from Hox clusters of vertebrates, myriapods and annelids can contain the conservative parts in the zones overlapping with proteincoding sequences, especially if this sequence is homeobox. However, this does not imply a structural or functional conservatism of these lncRNAs. It is more likely that bilateral animals with intact Hox clusters use a similar principle of regulation, which mechanism is yet obscure.

We may expect animals with a disorganized or an atomized cluster to possess a poor repertoire of Hox-associated lncRNAs. For example, the topological homologue of *Hotairm1* was not found in the ascidian *Ciona intestinalis* although it exists in tetrapods, teleosts and even in amphioxus (Herrera-Úbeda et al., 2019). In Drosophila, whose cluster is disorganized, although to a lesser extent, the number of Hox lncRNAs is significantly reduced compared to vertebrates. Unfortunately, this hypothesis remains a raw guess, because the data obtained on the model objects are patently insufficient for testing it.

## Conclusion

Bilateral animals possess a rich repertoire of variable mechanisms for controlling the functioning of Hox clusters. Many of these mechanisms are lineage-specific, such as the system of cis-regulatory modules, which integrate the incoming signals from Gap proteins and segmentation proteins during insect development. Mammals use lincRNAs (HOTAIR, HOTTIP, etc.), which are absent in protostomes. However, both insects and vertebrates possess Hox regulation by single specific enhancers or intergenic lncRNAs, which implies the plesiomorphy of this regulatory principle. We suggest that NATs of Hox genes found in mammals, myriapods and annelids are an example of such an ancestral regulatory mechanism, with individual elements having evolved independently in different bilaterian lineages. Hox NATs system is still poorly understood, and the nereidid annelids seem to be a good model for studies of this kind. In these animals NATs of almost all Hox genes can be easily found and analyzed. Further comparative analysis of Hox NATs functioning in the representatives of three evolutionary distinct clades is needed for the reconstruction of the regulatory evolution of Hox clusters.

## Materials and methods

### Animals

Adult *Alitta virens* were collected in the Chupa Inlet, the White Sea, near the “Kartesh” marine research station of the Zoological Institute of the Russian Academy of Sciences. Mature animals were caught with a hand net near the water surface during the spawning period (June and July). Artificial fertilization and cultivation of the embryos were carried out at 10.5°C. The culture of postlarval animals was kept in the laboratory of experimental embryology (Petergof, Russia) under the following conditions: temperature, 18°C; salinity, 23^0^/_00_; artificial sea water (Red Sea salt). The culture of *Platynereis dumerilii* is kept in the laboratory of experimental embryology at the same temperature, salinity 23 artificial sea water (Red Sea salt).

### Regeneration experiment

Juvenile worms consisting of 20-30 segments (*A. virens)* and 15-20 segments (*P. dumerilii)* were relaxed in clove oil (Sigma) of a low concentration for 5 min and cut into two pieces. Experimental details are provided in Novikova et al., 2013.

### Gene cloning

Gene cloning was performed in the Laboratory for Development and Evolution, Department of Zoology, University of Cambridge. Total RNA was isolated from pygidia and several posterior segments of young regenerates (7-10 days of regeneration) of *Alitta virens* and *Platynereis dumerilii* using TRIzol reagent (Invitrogen), according to the manufacturer’s instructions. To obtain 3’ and 5’ ends of transcripts of *Avi-antiHox4, Avi-antiHox5, Pdum-Lox4, Pdum-Lox2* and *Avi-Post2* the SMART™ (Switching Mechanism At 5’ end of the RNA Transcript) RACE method (SMART™ RACE cDNA amplification kit, Clontech) was used (Brena et al., 2006). To design RACE primers, the sequences of Hox genes of *A. virens* and *P.dumerilii* described earlier were used (Andreeva et al., 2001). Several forward and reverse primers for each Hox sequence were generated. To obtain 3’-ends of antisense (as) transcripts and 5’-ends of sense transcripts, reverse primers were used. 5’-ends of antisense transcripts and 3’-ends of sense transcripts were obtained with forward primers. In this paper, we denote the described or mentioned sequences with the help of abbreviations currently used in GenBank for the studied animals: *Avi* for *Alitta virens* and *Pdum* for *Platynereis dumerilii*. However, many genes of these animals, including a few antisense ncRNAs described in this work, are registered in GenBank under old designations: *Nvi* for *Alitta virens* (from the former species name *Nereis virens*) and *Pdu* for *Platynereis dumerilii*. GenBank accession numbers for the cDNA sequences are as follows: *Avi-antiHox4_1* - KX998894.1; *Avi-antiHox4_2* - KX998895.1; *Avi-antiHox5* - KP100547.1.

### Whole mount in situ hybridization (WMISH) and double WMISH

We used similar WMISH protocols for *A. virens* and for *P dumerilii.* For details of WMISH of larval stages see Kulakova et al., 2007, of postlarval stages, see Bakalenko et al., 2013, of regeneration, see Novikova et al., 2013. Minor modifications applied to the previously developed WMISH protocols for the detection of antisense transcripts were as follows: increasing the time and the number of washings after incubation with the probe and the antibodies, increasing the temperature of washings up to 68° C after the probe incubation, increasing the time of BM-Purple (Roche) incubation up to 48 hours. The detailed protocols are available upon request. Double WMISH was performed using two substrates: BM-Purple and Fast Red RC (Sigma Aldrich). For double WMISH probe synthesis, we used DIG RNA Labeling Mix (for BM-Purple detection) and Fluorescein RNA Labeling Mix (for Fast Red detection).

## Acknowledgements

The authors are grateful to Professor Michael Akam for providing the opportunity to N.B. to perform cloning in his Laboratory for Development and Evolution, Department of Zoology, University of Cambridge and to Carlo Brena for priceless methodical recommendations and support in cloning Hox antisense transcripts. We thank the staff of the White Sea Biological Station “Kartesh” (Zoological Institute, Russian Academy of Science) for help in collecting and maintaining *A. virens.* The research was performed using the facilities of the Research park of Saint Petersburg State University “CHROMAS” and “Culture Collection of Microorganisms”.

## Author Contributions

N.I.B. performed cloning. E.L.N., M.A.K. and N.I.B. performed WMISH. E.L.N. and M.A.K. conceptualized the project, analyzed the results, interpreted the results and wrote the manuscript. M.A.K. supervised the project. E.L.N. performed regeneration experiments and acquired funding (18-04-00450), M.A.K. participated in grant research (19-14-00346).

