## Supplementary Table S1, Table S2, Fig.S1. for "Annelids win again: the first evidence of Hox antisense transcription in Spiralia"

*Corresponding authors:

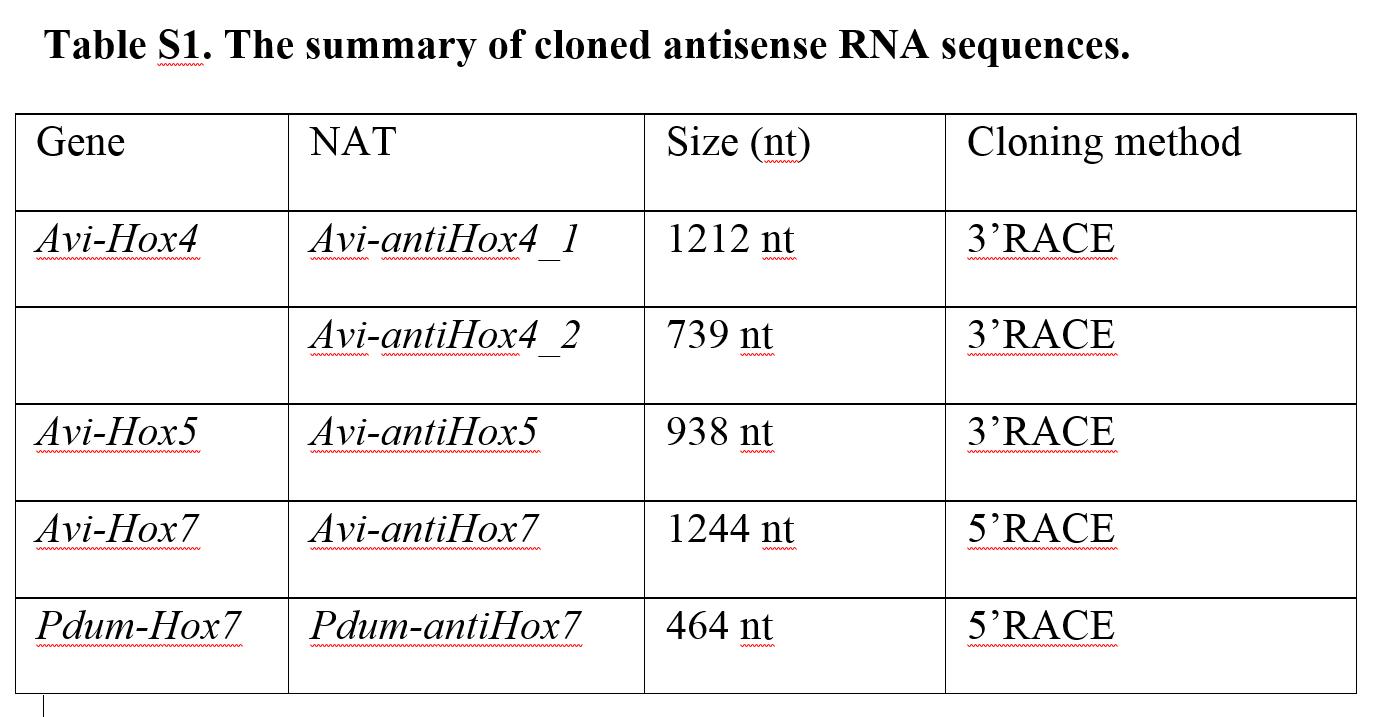

**Table S1. The summary of cloned antisense RNA sequences.**

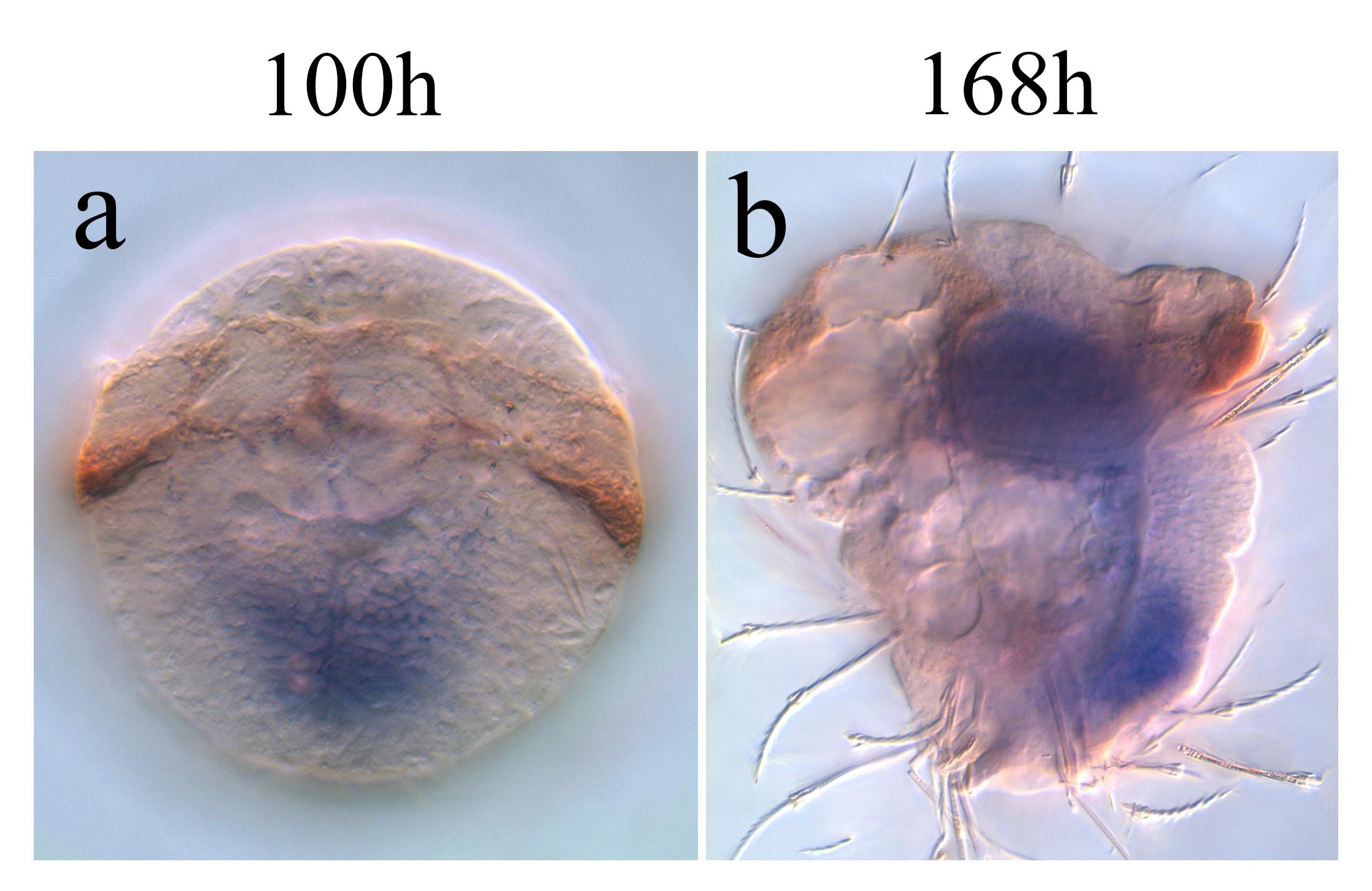

**Fig. S1. Antisense transcription of *Hox7* gene in larvae of *A. virens.*** On (**a**) and (**b**) the late trochophore (ventral view) and late metatrochophore (lateral view) are shown respectively.

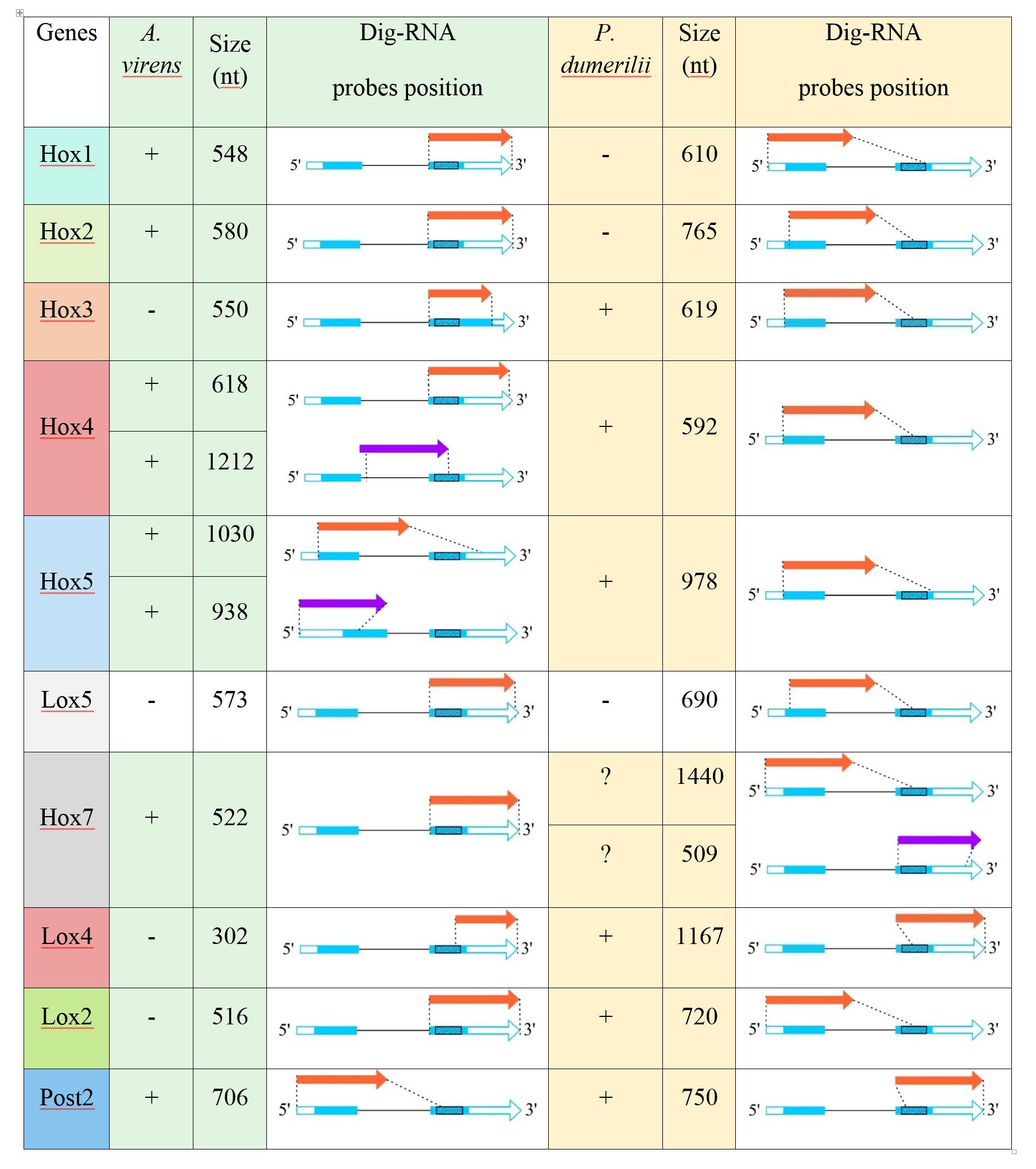

**Table S2. The summary of analyzed antisense lncRNAs of *A. virens* and P. *dumerilii****.* Red arrows indicate the position of sense probes. The purple color indicates the position of antisense probes to cloned lncRNA sequences. Blue boxes indicate protein-coding sequences. White boxes indicate 3’- and 5’-non-coding areas. Black shaded box indicates the position of the homeobox.
